# G_4_C_2_ repeat RNA mediates the disassembly of the nuclear pore complex in C9orf72 ALS/FTD

**DOI:** 10.1101/2020.02.13.947721

**Authors:** Alyssa N Coyne, Benjamin L Zaepfel, Lindsey Hayes, Boris Fitchman, Yuval Salzberg, Kelly Bowen, Hannah Trost, Frank Rigo, Amnon Harel, Clive N Svendsen, Dhruv Sareen, Jeffrey D Rothstein

## Abstract

Nucleocytoplasmic transport, controlled by the nuclear pore complex, has recently emerged as a pathomechanism underlying neurodegenerative diseases including C9orf72 ALS/FTD. However, little is known about the underlying molecular events and the underlying biology in human neurons. Using super resolution structured illumination microscopy of twenty three nucleoporins in nuclei from C9orf72 iPSC derived neurons and postmortem human tissue we identify a unique subset of eight nucleoporins lost from human neuronal nuclei. POM121, an integral transmembrane nucleoporin, appears to coordinate the composition of the nucleoporins within human neuronal nuclei ultimately impacting nucleocytoplasmic transport, and subsequent cellular toxicity in C9orf72 iPSNs. These data suggest that POM121 is a critical nucleoporin in the maintenance of the nuclear localization of specific nucleoporins in human neurons. Moreover, loss of nuclear POM121, as a result of expanded C9orf72 ALS/FTD repeat RNA, initiates a pathological cascade affecting nucleoporin composition within neuronal nuclei, nuclear pore complex function, and overall downstream neuronal survival.

## Introduction

The most common genetic cause of familial and sporadic forms of the motor neuron disease Amyotrophic Lateral Sclerosis (ALS) is a GGGGCC (G_4_C_2_) hexanucleotide repeat expansion (HRE) in intron 1 of the *C9orf72* gene (DeJesus-Hernandez et al., 2011; Renton et al., 2011). Notably, the *C9orf72* HRE is also causative of the second most common form of dementia, Frontotemporal dementia (FTD) affecting neurons within the frontal and temporal cortices (Ling et al., 2013). The HRE is bidirectionally transcribed to form sense (G_4_C_2_) and antisense (G_2_C_4_) RNA species that pathologically accumulate into nuclear RNA foci. Repeat associated non-ATG translation (RANT) of G_4_C_2_ and G_2_C_4_ RNA produces five dipeptide repeat (DPR) proteins [poly(GA), poly(GP), poly(GR), poly(PR), poly(PA)]. Together, these RNA species and DPR proteins are thought to contribute to disease through gain of toxicity mechanisms. In addition, the *C9orf72* HRE may also lead to haploinsufficiency of C9ORF72 protein resulting in a mild cellular, but not clinical, loss of function phenotype (Gitler and Tsuiji, 2016; Taylor et al., 2016). Recently, alterations is an essential cellular process, nucleocytoplasmic transport (NCT), have been identified in models of C9orf72 ALS/FTD, including human, rodent, fly, and yeast (Freibaum et al., 2015; Jovicic et al., 2015; Zhang et al., 2015). However, the molecular mechanisms underlying these disruptions and the normal biology governing these cellular processes in human neurons remain largely unknown.

The mammalian nuclear pore complex (NPC) is a ∼120 MDa, ∼450 protein complex consisting of multiple copies of approximately 30 nucleoporins highly organized with eight-fold rotational symmetry that can be segregated into four domains of the nuclear pore: cytoplasmic ring and filaments, central channel, scaffold, and nuclear basket. The domains can be further subdivided into nucleoporin subcomplexes such as the Y complex, transmembrane nucleoporins, the Nup62 complex, and the cytoplasmic and nuclear basket complexes, all of which have defined roles in nuclear pore assembly, maintenance and function (Lin and Hoelz, 2019; Raices and D’Angelo, 2012). The NPC is a barrier between the nucleus and cytoplasm and controls NCT, chromatin organization and gene expression (Beck and Hurt, 2017; Raices and D’Angelo, 2012, 2017). About one third of nucleoporins (Nups) contain FG repeat domains that are integral components of the central channel, cytoplasmic filaments, and nuclear basket. These intrinsically disordered domains interact with nuclear transport proteins and form a selective permeability barrier to control the transport of macromolecules (Li et al., 2016). A subset of these FG rich Nups can also have additional NPC independent functions in regulating gene transcription (Raices and D’Angelo, 2012). On the other hand, scaffolding Nups, such as the transmembrane and Y complex Nups, coordinate the formation of nuclear pores and anchor the pore in the nuclear envelope (Beck and Hurt, 2017; Harel et al., 2003; Mitchell et al., 2010).

The NPC critically maintains passive and active NCT. While macromolecules less than 40-60 kDa can passively diffuse through the nuclear pore, those greater than 40-60 kDa must be actively transported. The process of active transport relies on interactions between importins, exportins, Ran GTPase (Ran), and proper maintenance of the Ran gradient. In the nucleus, RCC1 mediates the conversion of Ran GDP to Ran GTP, and in the cytoplasm, RanGap1 facilitates the hydrolysis of Ran GTP to Ran GDP resulting in Ran GTP residing primarily in the nucleus and Ran GDP in the cytoplasm (Melchior, 2001; Raices and D’Angelo, 2012). Thus, the localization of Ran has been considered a static assessment of NCT (Eftekharzadeh et al., 2018; Grima et al., 2017; Zhang et al., 2015).

Most Nups are extremely long-lived in non-dividing cells including neurons, many having half-lives measured in months to years. As a result, alterations in NPCs and NCT have been reported during aging (D’Angelo et al., 2009; Hetzer, 2010; Savas et al., 2012; Toyama et al., 2013). However, little is actually known about the structure and overall biology of the NPC in human neurons, especially in the context of neural injury and disease. While mutations in specific Nups (e.g Nup62 and Aladin) or nuclear pore associated proteins, such as the nuclear transport protein Gle1, have been linked to neuronal degeneration (Nofrini et al., 2016; Sakuma and D’Angelo, 2017), nothing is known about the actual alterations of NPC structure and function in these rare diseases. In age-related neurodegenerative diseases such as ALS, Huntington’s Disease (HD), and Alzheimer’s Disease (AD), disruptions in NCT appear to be exacerbated over time (Chou et al., 2018; Eftekharzadeh et al., 2018; Freibaum et al., 2015; Gasset-Rosa et al., 2017; Grima et al., 2017; Jovicic et al., 2015; Zhang et al., 2015). In addition, specific Nups have been reported as modifiers of C9orf72 mediated toxicity in the *Drosophila* eye (Freibaum et al., 2015) and mislocalization of the nuclear pore associated protein RanGAP1 and Nups has been reported in postmortem human tissue and multiple mouse models based on overexpression of *C9orf72* or individual DPRs (Chew et al., 2019; Zhang et al., 2015; Zhang et al., 2018b; Zhang et al., 2016; Zhang et al., 2019b). Collectively, these studies suggest that altered NCT is a primary pathological feature of ALS and FTD. However, the normal biological role of specific Nups in human neurons, the precise disruptions in the nuclear expression and localization of NPC components, their impact on NCT, and any potential biological links to cellular injury in neurons harboring the endogenous *C9orf72* repeat expansion remain unknown. In order to mechanistically understand how alterations in the NPC and NCT contribute to disease pathogenesis, a comprehensive evaluation of the precise deficits and their role in human neuronal biology is essential.

Here, using an induced pluripotent stem cell (iPSC) derived neuron (iPSN) model of *C9orf72* ALS and super resolution structured illumination microscopy (SIM), we show that a specific and reproducible subset of eight Nups are lost from human neuronal nuclei. Notably, these alterations are closely mimicked in actual postmortem human motor cortex and thoracic spinal cord. Mechanistically, genetic manipulation of candidate altered Nups reveals that the transmembrane Nup POM121 plays an integral role in maintaining the nuclear composition of eight specific NPC components in iPSNs. Loss of POM121 in C9orf72 ALS/FTD is not mediated by DPRs or loss of C9ORF72 protein but occurs in the presence of pathologic G_4_C_2_ repeat RNA. These data provide direct evidence that pathologic repeat RNA initiates early disruptions to the nuclear pore complex in *C9orf72* mediated disease. Ultimately, alterations in NPC associated POM121 subsequently impacts the nuclear repertoire of specific Nups and together these alterations impact the distribution of Ran GTPase and cellular toxicity. These data suggest that POM121 mediated maintenance of nucleoporin composition within human neuronal nuclei plays an essential role in the function of NPCs and overall human neuronal survival. Together, our data highlight the NPC and POM121 as a potential therapeutic target in *C9orf72* ALS.

## Results

### Characterization of *C9orf72* pathology in an accelerated iPSC derived spinal neuron differentiation protocol

While the use of postmortem human tissue is valuable for identification of pathological hallmarks of disease, the study of neurodegenerative diseases such as ALS through use of iPSCs provides a unique and essential opportunity to identify underlying molecular mechanisms of disease pathogenesis. Neurons derived from human iPSCs maintain endogenous expression of disease associated genes and proteins rather than relying on overexpression in non-human animal models. However, many previous iPSC differentiation protocols take months to generate spinal neurons leading to increased differentiation and batch variability (Sances et al., 2016; Zhang et al., 2019a). To model *C9orf72* ALS with iPSCs, spinal neurons were differentiated using the direct induced motor neuron (diMNs) protocol (**Fig. S1A**) in which terminal differentiation and neuronal maturation begins at day 12 of differentiation with the addition of specific neuronal maturation growth factors (see **Experimental Procedures**, **Fig. S1A**). Ultimately, a reproducible population of spinal neurons is generated by day 18 of differentiation of which about 30% of which are Islet-1 positive lower motor neurons (**Fig. S1A-B**). Importantly, *C9orf72* ALS/FTD clinically and pathologically affects multiple different populations of neurons including interneurons, upper motor neurons, and neurons within the frontal and temporal cortices across two clinically distinct but genetically overlapping neurodegenerative diseases ALS and FTD (Cook and Petrucelli, 2019; Ferrari et al., 2011). As more than just Islet-1 positive lower motor neurons are affected in disease, the analysis of all neurons within iPSC derived spinal neurons cultures is essential for a comprehensive understanding of pathogenic mechanisms. As a result, we began by evaluating *C9orf72* pathology in these mixed iPSN cultures. Similar to our previous publications showing iPSNs produce G_4_C_2_ repeat RNA which accumulate into RNA foci (Donnelly et al., 2013), qRT-PCR experiments reveal an age dependent increase in both G_4_C_2_ and G_2_C_4_ repeat RNA from day 18 to day 32 of differentiation (**Fig. S1C**) suggesting either increased production or accumulation of repeat RNA. Conversely, Poly(GP) DPR levels, as detected by immunoassay, remain constant over time for each individual iPSC line tested (**Fig. S1D**) consistent with our previous studies in human CSF showing variability in absolute Poly(GP) levels amongst individual patients (Gendron et al., 2017). Furthermore, we find a slight reduction in *C9orf72* RNA levels (**Fig. S1E**) and no change in C9ORF72 protein (lower band as determined by western blot on C9orf72^-/-^ iPSC line (data not shown; **Fig. S1F-G**)) at day 32 of differentiation. Taken together, these data suggest that while our iPSN model of *C9orf72* ALS recapitulates pathologic repeat RNA and DPR production, it does not result in haploinsufficiency of C9ORF72, consistent with variability in C9ORF72 protein levels seen in actual patients (Sareen et al., 2013; Waite et al., 2014).

A major pathologic hallmark of the large majority of ALS and FTD cases, including *C9orf72*, includes the cytoplasmic mislocalization and accompanying accumulation and aggregation of the normally nuclear RNA binding protein TDP-43 (Ling et al., 2013). Using immunostaining and confocal imaging, we show that TDP-43 does not mislocalize or accumulate in the cytoplasm of *C9orf72* iPSNs compared to controls (**Fig. S1H-I**) at day 32 of differentiation. In agreement with human biopsy and autopsy studies (Vatsavayai et al., 2016), this suggests that TDP-43 mislocalization may be a much later event in ALS pathogenesis. In contrast, while we see no overt difference in cell death at baseline, *C9orf72* iPSNs are susceptible to glutamate induced excitotoxicity (**Fig. S1J**) similar to our previous reports using a much longer spinal neuron differentiation protocol (Zhang et al., 2015).

### Specific nucleoporins are altered in *C9orf72* iPSN nuclei in an age dependent manner

We have previously reported that the NPC associated protein RanGAP1 can be sequestered by G_4_C_2_ repeat RNA to impact functional NCT (Zhang et al., 2015). Given that the NPC critically governs NCT (Lin and Hoelz, 2019; Raices and D’Angelo, 2012), we hypothesized that specific alterations in the individual Nup building blocks of the NPC may also be a pathological consequence of the *C9orf72* HRE. To examine individual Nups within neuronal nuclei, we used super resolution structured illumination microscopy (SIM) which allows for increased resolution of Nup immunoreactivity albeit not at the resolution of individual Nup molecules (Schermelleh et al., 2008). We evaluated the nuclear localization and expression of 23 of the 30 human Nups in NeuN positive nuclei isolated from iPSNs at both day 18 and day 32 of differentiation, in up to 9 different *C9orf72* iPSC lines and 11 different control iPSC lines (see **Table S1**). Following SIM, we used automated analytics to determine the number of Nup spots. Although this approach was employed for most Nups, for a few Nups individual spot analytics was not possible due to limits of resolution in distinguishing individual spots possibly due to their abundance or distribution throughout the nucleus and nuclear membrane. Furthermore, the resolution achieved with SIM is often dependent on the fluorophore used for immunostaining (Cox, 2015; Wegel et al., 2016). In cases where individual spots could not be resolved, we calculated the percent of the total nucleus volume occupied by the Nup (see **Experimental Procedures** for additional details). In addition to variable resolution amongst individual Nups, we note that potential differences in the number of Nup molecules per pore (e.g. multiples of 8) and differential antibody affinities does not allow for a comparison of absolute Nup spot or volume numbers amongst each of the 23 Nups evaluated. Thus, all analyses were conducted on an individual Nup basis comparing *C9orf72* to controls. Using this approach, we reproducibly found that 8 of the 23 Nups examined including the nuclear basket Nups Nup50 and TPR, the central channel Nup Nup98, all three transmembrane Nups GP210, NDC1, and POM12**1**, and the Y complex outer ring Nups Nup107 and Nup133 are lost from *C9orf72* iPSN nuclei at day 32 but not day 18 when compared to controls at the respective time point (**Fig. 1**). These data highlight the age dependence of specific Nup alterations in the maturation of pathogenic cascades in *C9orf72* ALS. Importantly, we observe a loss of nuclear POM121 in Islet1 positive lower motor neurons at day 32, but not at day 18 (**Fig. S2A-B**) and in spinal neurons derived using a previously described iPSN differentiation protocol (Donnelly et al., 2013; Zhang et al., 2015) (**Fig. S2C-D**). Together, these data suggest that our observed alterations in specific Nups are reproducible in multiple neuronal subtypes affected in ALS pathogenesis and across multiple differentiation protocols. Notably, nuclear loss of POM121 was confirmed with a second antibody recognizing the N-terminal, nuclear envelope associated region of the protein (**Fig. S3A-B**). While isolated nuclei preparations provide for more accurate reconstruction and increased resolution of Nup immunostaining within the nuclear membrane and nuclei, we observe a similar decrease in POM121 nuclear intensity using standard confocal microscopy in intact iPSNs (**Fig. S3C-D**) suggesting that our results are not an artifact of SIM or the process of isolating nuclei. Furthermore, we do not observe mislocalization of POM121 from the nucleus to the cytoplasm in intact iPSNs (**Fig. S3C**). Moreover, western blots quantitatively confirm a decrease in total POM121 protein in isolated nuclei from *C9orf72* iPSNs (**Fig. S3E-F**) suggesting that our results are not an artifact of imaging paradigms. Additionally, decreased nuclear expression and localization of POM121 and other affected Nups is not a result of decreased mRNA levels (**Fig. S3G**). Together, these data suggest that post-transcriptional alterations result in decreased nuclear localization of specific Nups without corresponding cytoplasmic mislocalization or accumulation. In spite of the loss of these 8 specific nucleoporins from the nucleus, we do not find alterations in overall nuclear pore number or distribution of pores based on immunostaining with a general nuclear pore antibody that recognizes 4 FG repeat containing Nups, Nup153, Nup62, Nup214, and Nup358/RanBP2 (mAb414; 414) (**Fig. S4A-C**).

**Figure 1:**
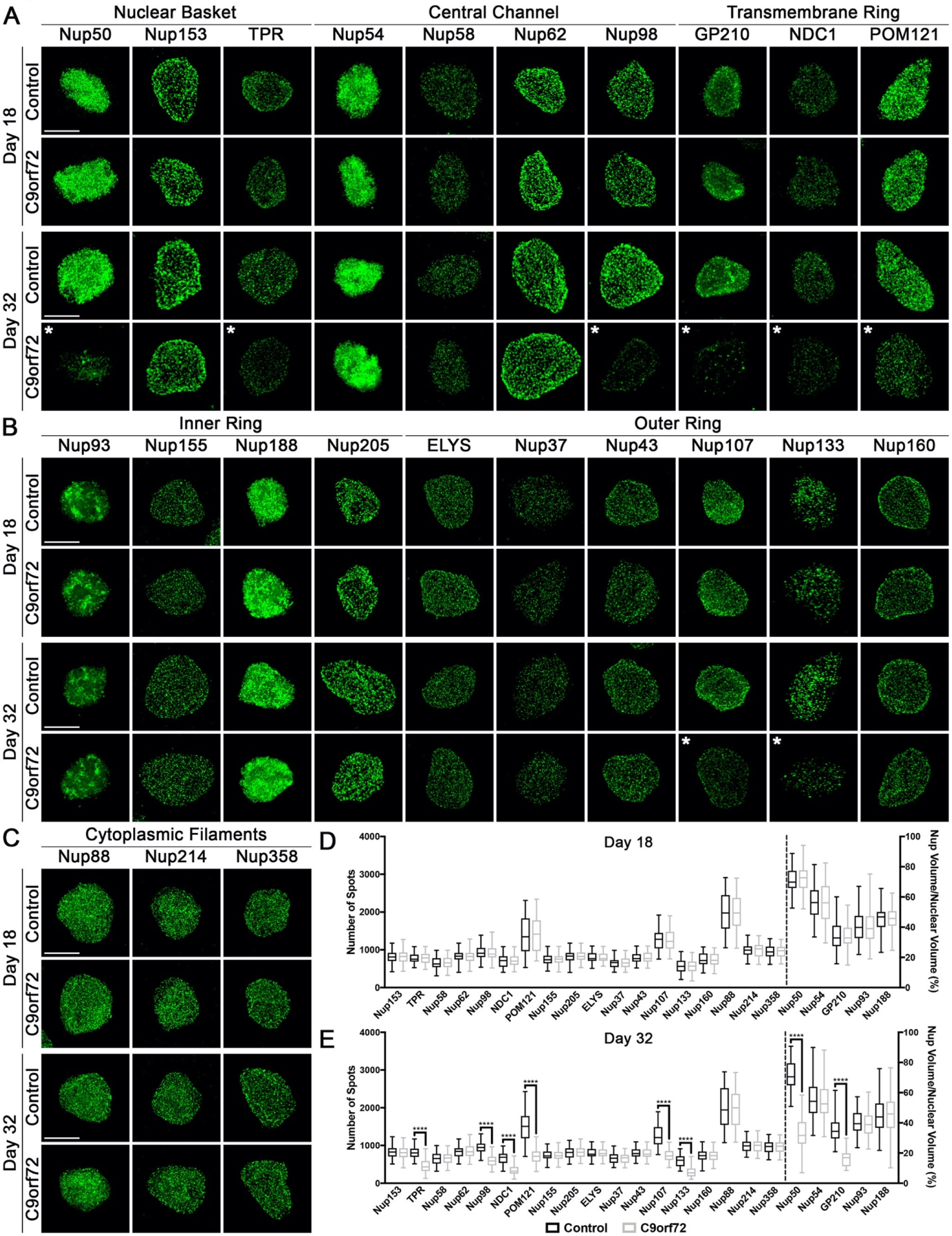
Specific nucleoporins are altered in nuclei from *C9orf72* iPSNs. (**A-C**) Maximum intensity projections from SIM imaging of Nups in nuclei isolated from control and *C9orf72* iPSNs. Genotype and time point as indicated on left, antibodies as indicated on top. (**D-E**) Quantification of Nup spots and volume. Time point as indicated on top. N = 8 control and 8 *C9orf72* iPSC lines (including 1 isogenic pair), 50 NeuN+ nuclei per line/time point. Two-way ANOVA with Tukey’s multiple comparison test was used to calculate statistical significance. **** p < 0.0001. Scale bar = 5 μm. * indicates significantly altered Nups.

To directly compare control and *C9orf72* iPSN nuclear envelopes at high resolution, we used nuclear exposure and direct surface imaging by field emission scanning electron microscopy (SEM). Here we found that that the overall number and distribution of NPCs is unchanged in *C9orf72* iPSN nuclei (**Fig. S4D**). Furthermore, higher magnification imaging focusing on individual NPCs shows no overt change in NPC architecture in *C9orf72* IPSN nuclei (**Fig. S4D**). Together, these data suggest that individual Nup molecules may be missing from the nuclear envelope, presumably from pore octamers, without overall loss of entire NPCs.

### *C9orf72* iPSN nuclei are not leaky

Given our previous report that active NCT is disrupted in *C9orf72* iPSNs (Zhang et al., 2015) and our data that the nuclear localization and expression of 8 Nups is reliably and reproducibly decreased in *C9orf72* iPSN nuclei without overall effects on NPC distribution and structure (**Fig. 1, Fig. S4**), we next wanted to determine if *C9orf72* iPSN nuclei were passively leaky. Normally, macromolecules less than 40-60 kDa can passively diffuse through the NPC and thus fluorescent dextrans of various sizes have frequently been used to evaluate the passive permeability of NPCs (Zhu et al., 2016). Confocal imaging of digitonin permeabilized iPSNs revealed that *C9orf72* nuclei are not passively leaky compared to controls (**Fig. S5**) suggesting that the overall passive permeability barrier of the NPC in intact despite the nuclear loss of 8 specific nucleoporins in *C9orf72* IPSNs.

### Nucleoporin alterations in postmortem *C9orf72* patient motor cortex and thoracic spinal cord are identical to those in *C9orf72* iPSNs

To determine whether our observed alterations in specific Nups in *C9orf72* iPSNs were reflective of real changes that occur in human patients or a consequence of *in vitro* differentiation and culturing of iPSNs, we next isolated nuclei from postmortem motor and occipital cortex from non-neurologic control and *C9orf72* patients and performed SIM. Of the 13 Nups analyzed, Nup50, TPR, Nup98, NDC1, POM121, Nup107, and Nup133 but not Nup153, Nup54, Nup62, Nup188, Nup205, or Nup160 are lost from NeuN positive nuclei from *C9orf72* patient motor cortex (**Fig. 2A-C**) but not occipital cortex (**Fig. 2A-B, D**), a brain region unaffected in ALS. Similar to our results in postmortem motor cortex, the nuclear expression and localization of 6 Nups (Nup50, TPR, NDC1, POM121, Nup107, and Nup133) are reduced in NeuN positive nuclei isolated from *C9orf72* patient thoracic spinal cord compared to controls (**Fig. S6**). Together, these data in patient tissue mimic our results in iPSNs. Notably, staining of postmortem paraffin embedded motor cortex tissue sections with an antibody recognizing the C-terminus of POM121 also shows a decrease in nuclear POM121 intensity (**Fig. S7A-B**), albeit without the resolution provided by SIM. Interestingly, co-staining of paraffin embedded tissue sections for POM121 and TDP-43 reveals that a reduction in POM121 intensity can be observed both in neurons with and without TDP-43 pathology (**Fig. S7C**). This suggests that that alterations to the nuclear composition of individual Nups and specifically the loss of POM121 may precede TDP-43 mislocalization which is thought to be a consequence of impaired NPC function. Together, these data suggest that our iPSN model recapitulates human disease pathogenesis, identifies specific nucleoporin alterations as a possible initiating pathologic event in *C9orf72* ALS, and highlights the use of our iPSN model as a tool for studying the mechanisms underlying disease alterations in individual NPC components.

**Figure 2.**
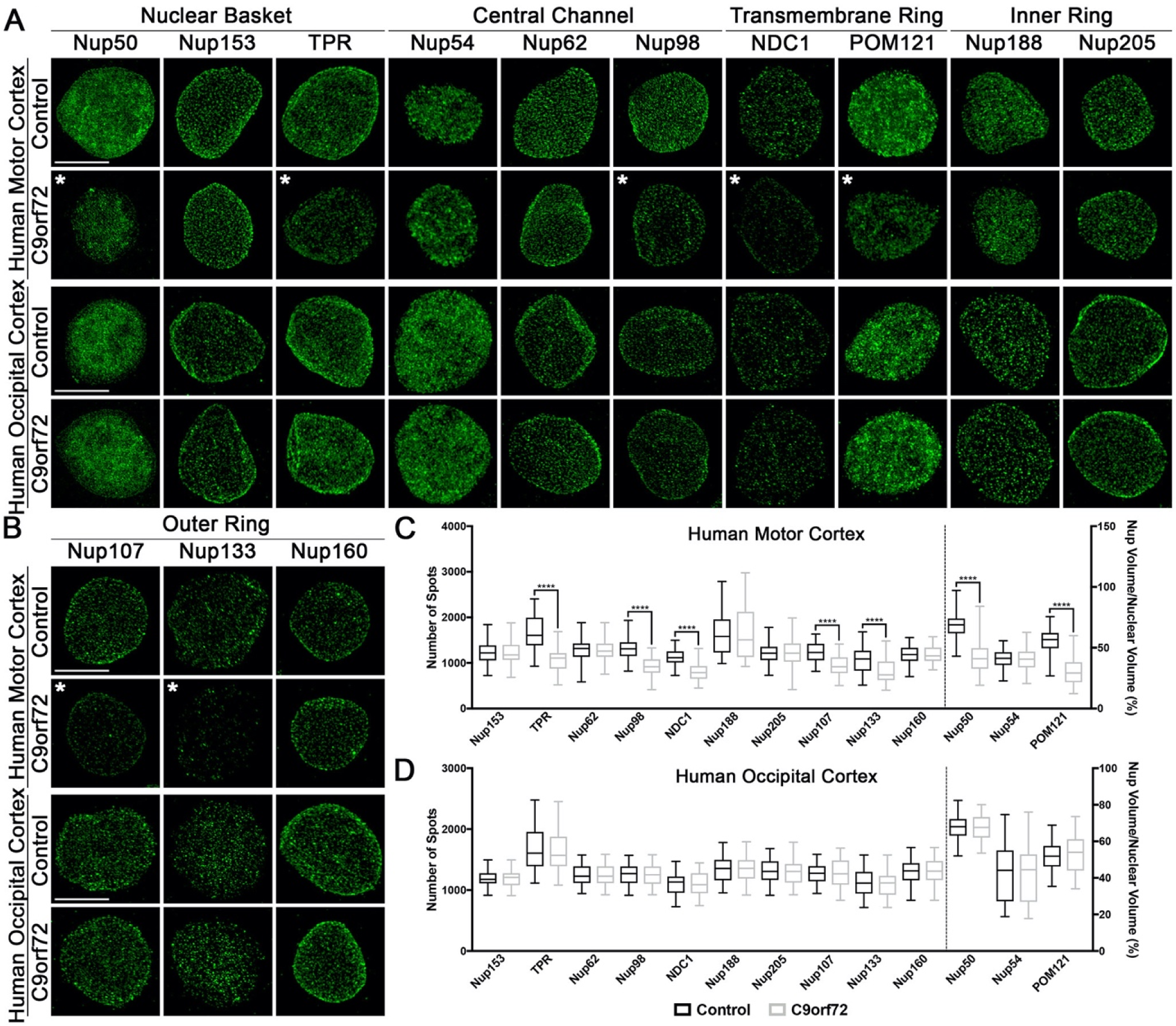
Specific nucleoporins are altered in nuclei from *C9orf72* patient motor cortex. (**A-B**) Maximum intensity projections from SIM imaging of Nups in nuclei isolated from postmortem human brain tissue. Genotype and brain region as indicated on left, antibodies as indicated on top. (**C-D**) Quantification of Nup spots and volume. Brain region as indicated on top. N = 3 control and 3 *C9orf72* cases, 50 NeuN+ nuclei per case. Two-way ANOVA with Tukey’s multiple comparison test was used to calculate statistical significance. **** p < 0.0001. Scale bar = 5 μm. * indicates significantly altered Nups.

### Overexpression of POM121 restores the nuclear expression and localization of specific Nups and Ran GTPase, mitigates alterations in functional NCT, and protects against cellular toxicity

The formation and intricate organization of the NPC is a highly coordinated process reliant on interactions amongst specific Nups and subcomplexes (Lin and Hoelz, 2019). While these relationships are highly studied in mitotic cells, little is known about their role in NPC maintenance in human neurons which are no longer dividing. To begin to understand the relationship between specific Nups in neurons and in disease, we overexpressed candidate Nups whose nuclear localization and expression is altered in *C9orf72* ALS iPSNs and used SIM to evaluate individual NPC components in isolated nuclei. To evaluate the ability of Nup98, Nup133, and POM121 to alleviate defects in nuclear composition of specific Nups, overexpression was conducted at a timepoint after the initial reproducible *C9orf72* mediated depletion of Nups from the nucleus (30 days, **Fig. 1**). While overexpression of the FG repeat containing central channel Nup, Nup98, only restored Nup98 nuclear localization and expression (**Fig. 3A-T**), overexpression of the structural Y complex Nup, Nup133, restored nuclear localization and expression of Nup50, TPR, Nup98, Nup107, and Nup133 but not the transmembrane Nups GP210, NDC1, and POM121 in *C9orf72* IPSNs (**Fig. 3A-T**). In contrast, overexpression of the transmembrane scaffolding Nup, POM121, at day 30 of differentiation, completely mitigated all alterations in nuclear expression and localization of Nups in *C9orf72* iPSNs within about 2 weeks, by day 46 of differentiation (**Fig. 3A-T**). Interestingly, Nup133 and POM121 overexpression does not alter the total number or distribution of NPCs as evaluated by immunostaining with the 414 antibody and SIM microscopy (**Fig. S8**). Together with our observations that there is no overall loss of NPC number or distribution in *C9orf72* iPSN nuclei, this suggests that overexpression POM121 may reassemble Nups into existing NPCs in human neurons. Conversely, a soluble variant of POM121 (sPOM121) which lacks the transmembrane domain and does not localize to the NPC (Franks et al., 2016), does not restore nuclear expression and localization of candidate Nups (**Fig. S9**). Furthermore, overexpression of the other transmembrane Nups, GP210 and NDC1, only restored the nuclear localization and expression of GP210 and NDC1 respectively (**Fig. S10**) and thus did not re-establish the nuclear composition of individual Nups to the extent of the transmembrane Nup POM121 (**Fig. 3A-T**). Interestingly, GP210, NDC1, and sPOM121 overexpression all partially restored the nuclear localization and expression of Nup98 (**Fig. S9A,C, Fig. S10A,D**) suggesting that Nup98 is highly sensitive to a variety of Nup manipulations. Together, these data suggest that the NPC associated variant of the transmembrane Nup POM121 is necessary for the re-establishment of proper nuclear localization and expression of specific Nups in *C9orf72* iPSNs.

**Figure 3:**
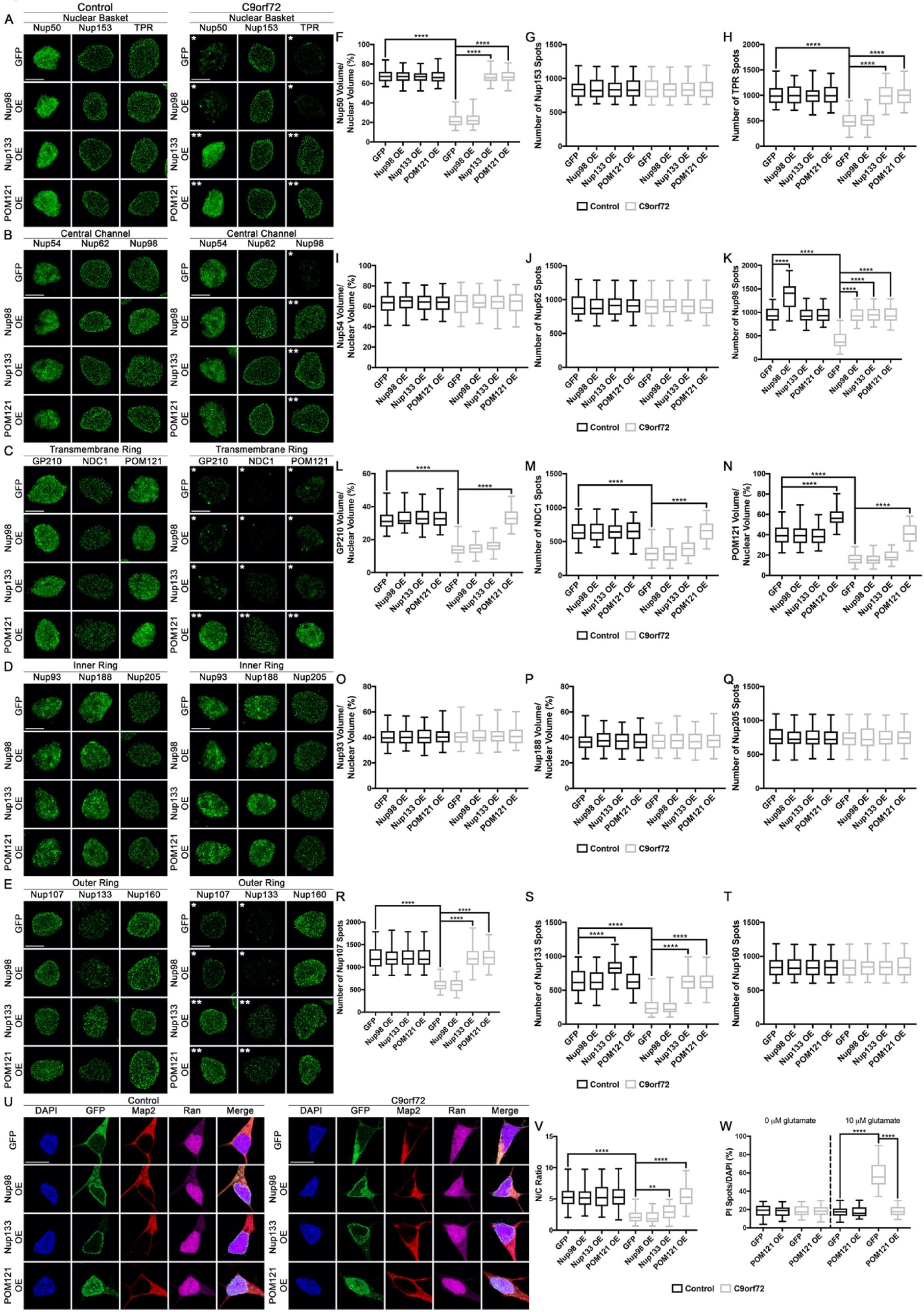
POM121 overexpression restores the nuclear expression and localization of specific Nups and Ran GTPase and mitigates cellular toxicity. (**A-E**) Maximum intensity projections from SIM imaging of Nups in nuclei isolated from control and *C9orf72* iPSNs overexpressing Nup98, Nup133, or POM121. Overexpression as indicated on left, genotype and antibodies as indicated on top. (**F-T**) Quantification of Nup spots and volume. N = 4 control and 4 *C9orf72* iPSC lines, 50 GFP+ nuclei per line/overexpression. Two-way ANOVA with Tukey’s multiple comparison test was used to calculate statistical significance. **** p < 0.0001. (**U**) Confocal imaging of control and *C9orf72* iPSNs overexpressing GFP tagged Nup98, Nup133, or POM121 immunostained for Ran. Overexpression as indicated on left, genotype and antibodies as indicated on top. (**V**) Quantification of nuclear to cytoplasmic ratio of Ran. N = 4 control and 4 *C9orf72* iPSC lines, at least 50 cells per line/overexpression. Two-way ANOVA with Tukey’s multiple comparison test was used to calculate statistical significance. ** p < 0.01, **** p < 0.0001. (**W**) Quantification of percent cell death following exposure to glutamate. N = 4 control and 4 *C9orf72* iPSC lines, 10 frames per well. Two-way ANOVA with Tukey’s multiple comparison test was used to calculate statistical significance. **** p < 0.0001. Scale bar = 5 μm (**A-E**), 10 μm (**U**). * indicates significantly altered Nups, ** indicates significantly restored Nups.

Given the integral role of the NPC in controlling and maintaining NCT and cellular function (Lin and Hoelz, 2019; Raices and D’Angelo, 2012), we next evaluated whether POM121 mediated compositional restoration of Nups within *C9orf72* iPSN nuclei affected the subcellular distribution of Ran GTPase, a critical component of the NCT machinery (Melchior, 2001; Raices and D’Angelo, 2012). Confocal imaging revealed that POM121 overexpression resulted in the relocalization of Ran from the cytoplasm to the nucleus in *C9orf72* iPSNs (**Fig. 3U-V**). Conversely, Nup98 overexpression had no effect on Ran localization and Nup133 overexpression only partially relocalized Ran to the nucleus (**Fig. 3U-V**). To evaluate whether POM121 overexpression could restore functional active NCT, we employed the NLS-tdTomato-NES (S-tdTomato) reporter construct previously described (Zhang et al., 2015) and monitored its subcellular localization over time following POM121 overexpression in control and *C9orf72* iPSNs (**Fig. S11A**). While S-tdTomato is observed to be predominately cytoplasmic by 3 days post transduction in *C9orf72* iPSNs, overexpression of POM121 completely restores the nuclear localization of the S-tdTomato reporter by 10 days post transduction (**Fig. S11B-E**). Together these data suggest that POM121 overexpression restores the localization of Ran GTPase and mitigates deficits in functional NCT in *C9orf72* iPSNs.

Next we asked whether POM121 mediated restoration of NPC composition and function was sufficient to mitigate the sensitivity of human neurons to exogenous stressors. Using glutamate induced cell death assays we find that in addition to its effects on nuclear expression and localization of Nups and functional NCT, POM121 overexpression suppressed glutamate induced excitotoxicity and promoted cellular survival in *C9orf72* iPSNs (**Fig. 3W, Fig. S12**). Together, these data suggest that POM121 overexpression can completely mitigate alterations in the nuclear expression and localization of specific Nups to restore NPC function and ultimately mitigate cellular toxicity.

### Loss of POM121 alone in wildtype neurons recapitulates disease phenotypes

Having established that restoring nuclear localization and expression of POM121 can re-establish the nuclear composition of specific Nups, repair NCT, and promote cellular survival, we sought to determine whether knock down of POM121 in wildtype iPSNs could recapitulate *C9orf72* mediated disease phenotypes. Due to the long half-life of many structural nucleoporins in neurons (Savas et al., 2012; Toyama et al., 2013), we employed the Trim Away method (Clift et al., 2017) to rapidly degrade endogenous Nup133, POM121, NDC1, or GAPDH protein in control iPSNs (**Fig. S13**). While knock down of Nup133 resulted in subsequent nuclear loss of Nup50, TPR, Nup107, and Nup133 (**Fig. 4A-P**), knock down of POM121 fully recapitulated the loss of Nup50, TPR, GP210, NDC1, POM121, Nup107, and Nup133 (**Fig. 4A-P**) from the nucleus as observed in *C9orf72* iPSNs (**Fig. 1**) and human motor cortex (**Fig. 2**) and thoracic spinal cord (**Fig. S6**) within 48 hours. Notably, this effect on the composition of Nups in iPSN nuclei was specific to the transmembrane nucleoporin POM121 as Trim21 mediated reduction of the transmembrane nucleoporin NDC1 had no effect on Nups other than itself (**Fig. 4A-P**). Interestingly, neither Nup133, NDC1, nor POM121 knock down resulted in disruptions in nuclear associated Nup98 (**Fig. 4A-P**). Furthermore, knock down of GAPDH, a non-NPC associated control protein, had no effect on nuclear expression and localization of any of the 15 Nups analyzed (**Fig. 4A-P**) suggesting that effects on nuclear expression and localization of Nups are specific to Nup manipulation itself. In addition to effects on the localization and expression of NPC components, Trim21 mediated knock down of POM121, but not Nup133 or GAPDH, resulted in Ran GTPase mislocalization from the nucleus to the cytoplasm (**Fig. 4Q-R**). Furthermore, Trim21 mediated knock down of POM121 increased the susceptibility of wildtype neurons to glutamate induced excitotoxicity (**Fig. 4S-T**). Together, these data suggest that nuclear loss of the transmembrane Nup, POM121 can initiate a pathological cascade reminiscent of *C9orf72* mediated disease alterations including disruption of NPC components and Ran GTPase as well as increased cellular sensitivity to stressors.

**Figure 4:**
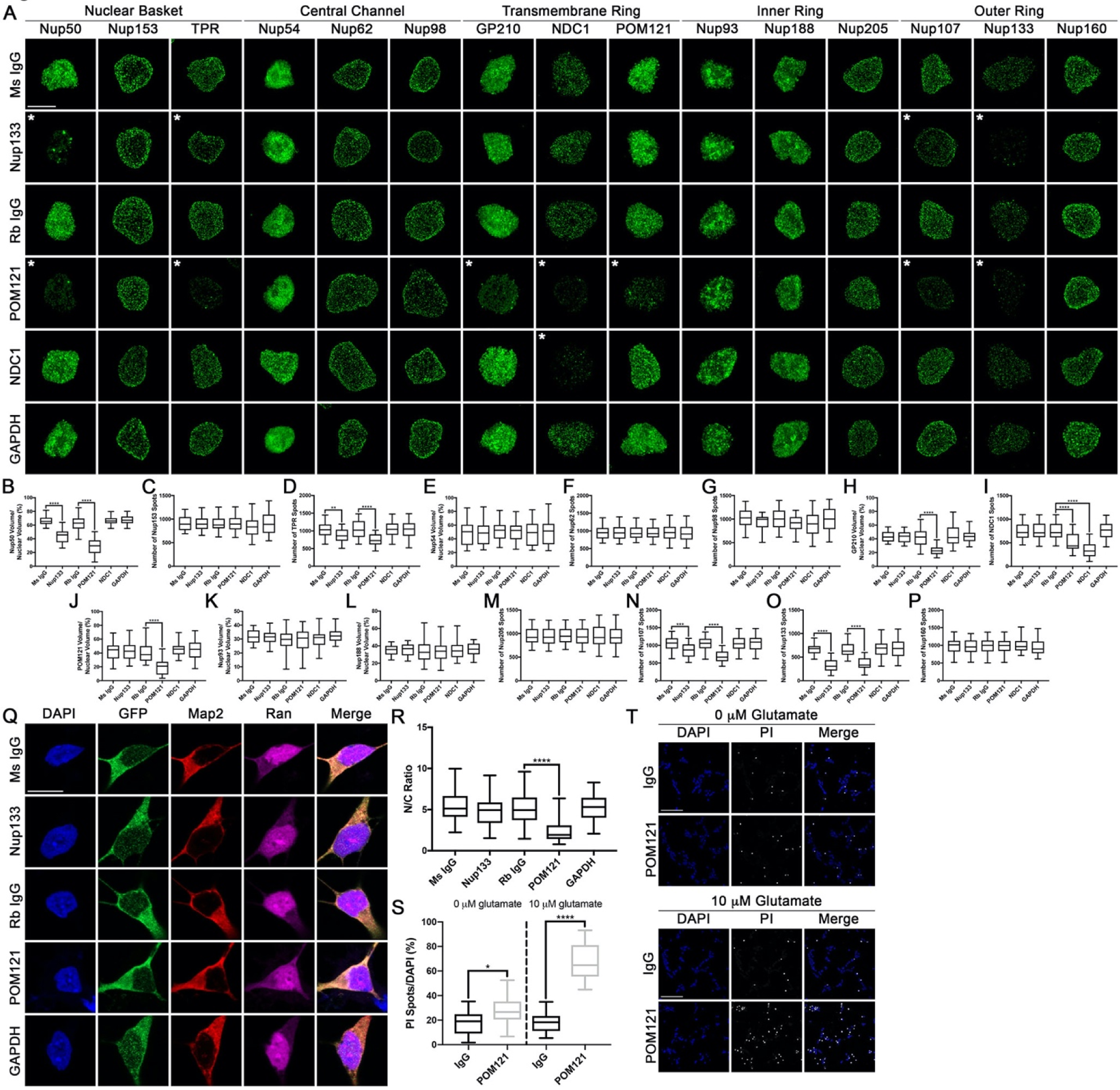
Reduction in POM121 in wildtype iPSNs recapitulates *C9orf72* mediated alterations in specific Nups and Ran GTPase and increases susceptibility to glutamate induced excitotoxicity. (**A-P**) Maximum intensity projections from SIM imaging (**A**) and quantification (**B-P**) of Nup spots and volume in nuclei isolated from wildtype iPSNs following knockdown of Nup133, POM121, NDC1, or GAPDH. Antibody used for Trim21 mediated knockdown as indicated on left, antibodies as indicated on top. N = 3 wildtype iPSC lines, 50 GFP+ nuclei per line/knockdown. One-way ANOVA with Tukey’s multiple comparison test was used to calculate statistical significance. ** p < 0.01, *** p < 0.001, **** p < 0.0001. (**Q**) Confocal imaging of control iPSNs following Trim21 GFP knockdown of Nup133, POM121, or GAPDH immunostained for Ran. Antibody used for Trim21 knockdown as indicated on left, antibodies as indicated on top. (**R**) Quantification of nuclear to cytoplasmic ratio of Ran. N = 3 wildtype iPSC lines, at least 50 cells per line/knockdown. One-way ANOVA with Tukey’s multiple comparison test was used to calculate statistical significance. **** p < 0.0001. (**S**) Quantification of percent cell death following exposure to glutamate. N = 3 control iPSC lines, 10 frames per well. Two-way ANOVA with Tukey’s multiple comparison test was used to calculate statistical significance. **** p < 0.0001. (**T**) Confocal imaging of cell death in control and C9orf72 iPSNs as measured by propidium iodide (PI) incorporation. Antibody used for Trim Away as indicated on left, glutamate concentration and stain as indicated on top. Scale bar = 5 μm (**A**), 10 μm (**Q**), 100 μm (**T**). * indicates significantly altered Nups.

### Loss of C9ORF72 and DPRs do not contribute to alterations in the POM121

It has been proposed that the *C9orf72* HRE exerts toxicity through the combination of three pathological phenomena including haploinsufficiency of C9ORF72 protein, accumulation of G_4_C_2_ and G_2_C_4_ repeat RNA, and production of toxic DPRs (Balendra and Isaacs, 2018; Cook and Petrucelli, 2019; Gitler and Tsuiji, 2016). Although we do not observe a reduction in C9ORF72 protein in our iPSN model (**Fig. S1F-G**), we first used a *C9orf72* knock out iPSC line to investigate whether complete loss of C9ORF72 protein affected the nuclear expression and localization of individual Nups. Using SIM, we found that none of the 22 Nups assessed were altered in nuclei isolated from *C9orf72*^-/-^ iPSNs compared to controls (**Fig. 5A-B**) suggesting that loss of C9ORF72 protein does not play a role in Nup alterations in *C9orf72* ALS.

**Figure 5:**
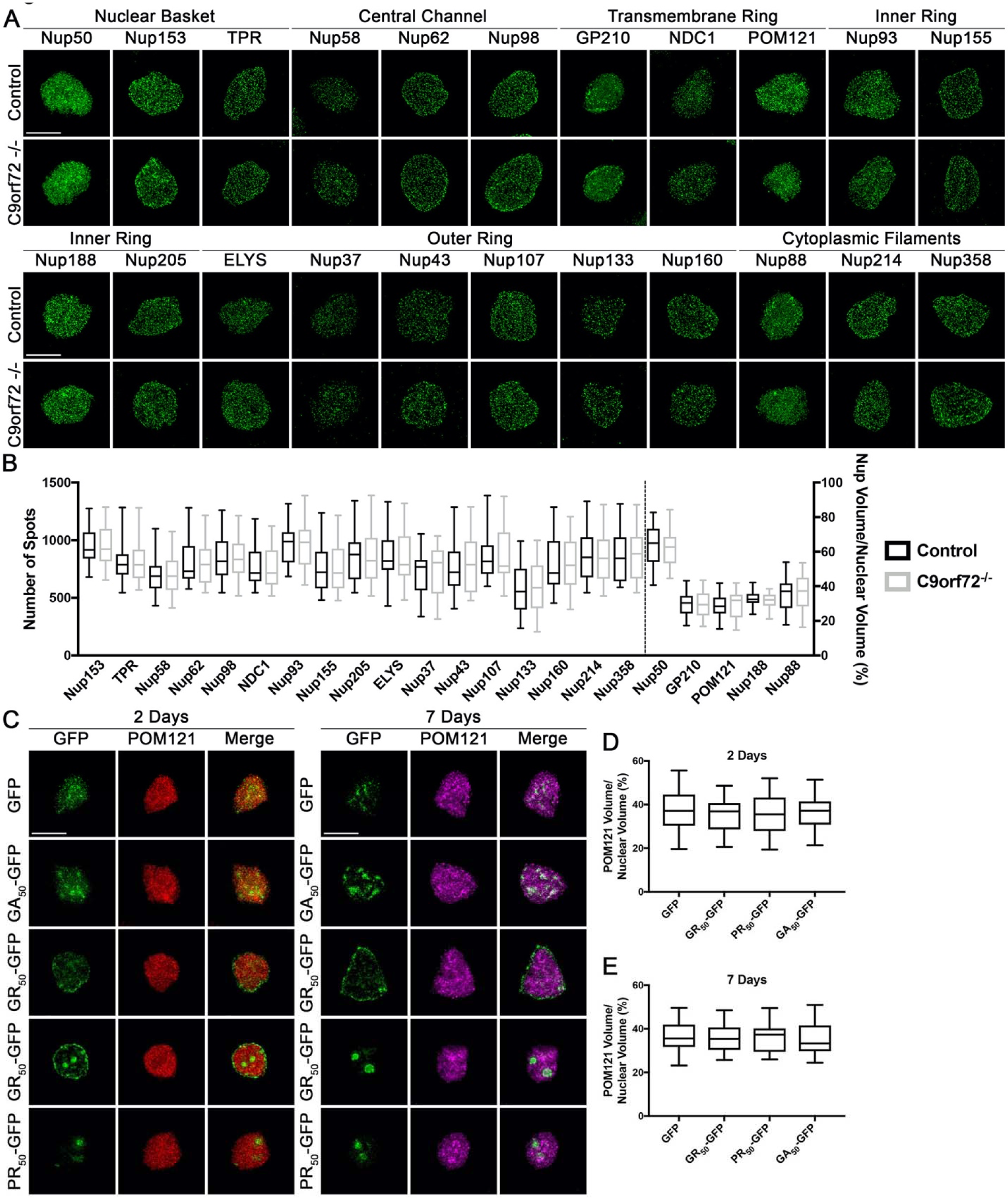
Loss of C9ORF72 protein or DPRs do not alter the nuclear localization and expression of POM121. (**A**) Maximum intensity projections from SIM imaging of Nups in nuclei isolated from control and *C9orf72^-/-^* iPSNs. Genotype as indicated on left, antibodies as indicated on top. (**B**) Quantification of Nup spots and volume. N = 1 control and 1 *C9orf72* null line, differentiations conducted in triplicate, 50 NeuN+ nuclei per line/differentiation. Two-way ANOVA with Tukey’s multiple comparison test was used to calculate statistical significance. (**C**) SIM imaging of POM121 in nuclei isolated from wildtype iPSNs overexpressing Poly(GA), Poly(GR), and Poly(PR) DPRs. Overexpression as indicated on left, antibodies and time points as indicated on top. (**D-E**) Quantification of percent total nuclear volume occupied by POM121 2 days (**D**) and 7 days (**E**) after overexpression of DPRs. N = 3 wildtype iPSC lines, 50 GFP+ nuclei per line/overexpression. One-way ANOVA with Tukey’s multiple comparison test was used to calculate statistical significance. Scale bar = 5 μm.

Multiple studies have suggested that artificial overexpression of DPRs produced by the *C9orf72* HRE can result in disruptions in NCT potentially via alterations including cytoplasmic mislocalization and aggregation of NPC associated proteins and Nups in mouse models, postmortem human tissue, and frog oocytes (Chew et al., 2019; Shi et al., 2017; Zhang et al., 2015; Zhang et al., 2018b; Zhang et al., 2016; Zhang et al., 2019b). While we do not observe cytoplasmic mislocalization or accumulation of POM121 in iPSNs (**Fig. S3C**) or human tissue (**Fig. S7**), we used SIM to evaluate whether overexpression of individual DPRs in wildtype iPSNs affected nuclear expression and localization of POM121. While both Poly(GR) and Poly(PR) strongly accumulate in and/or around the nucleus, nuclear localization of Poly(GA) was rarely observed (**Fig. 5C**). However, neither Poly(GA), Poly(GR), nor Poly(PR) affected the expression or distribution of POM121 within nuclei after 48 hours (**Fig. 5C-D**) or one week (**Fig. 5C,E**) of overexpression. These data indicate that like loss of C9ORF72 protein, DPRs are not responsible for the initiation of POM121 loss from human neuronal nuclei *C9orf72* disease pathogenesis. Furthermore, this suggests that repeat RNA itself may result in alterations in specific NPC components as was previously found to be the case for the NPC associated protein RanGAP1 (Zhang et al., 2015).

### Expression of G_4_C_2_ repeat RNA results in loss of POM121 from nuclei in iPSNs

To test whether pathological G_4_C_2_ RNA repeats were responsible for the reduced nuclear localization and expression of the integral transmembrane Nup POM121, we first treated control and *C9orf72* iPSNs with G_4_C_2_ targeting antisense oligonucleotides (ASOs) for only 5 days beginning at Day 30 of differentiation. After 5 day treatment with ASO, we observed a large reduction in G_4_C_2_ repeat RNA (**Fig. 6A**) but not the Poly(GP) DPR (**Fig. 6B**) indicating that this short ASO treatment paradigm can selectively reduce pathologic repeat RNA but not DPRs consistent with our previous reports that it takes about 3 weeks to completely reduce Poly(GP) levels in iPSNs (Gendron et al., 2017). Following nuclei isolation, we conducted SIM and found that this 5 day ASO treatment completely mitigated reduced nuclear localization and expression of POM121 in *C9orf72* IPSN nuclei compared to untreated and scrambled ASO controls (**Fig. 6C-D**).

**Figure 6.**
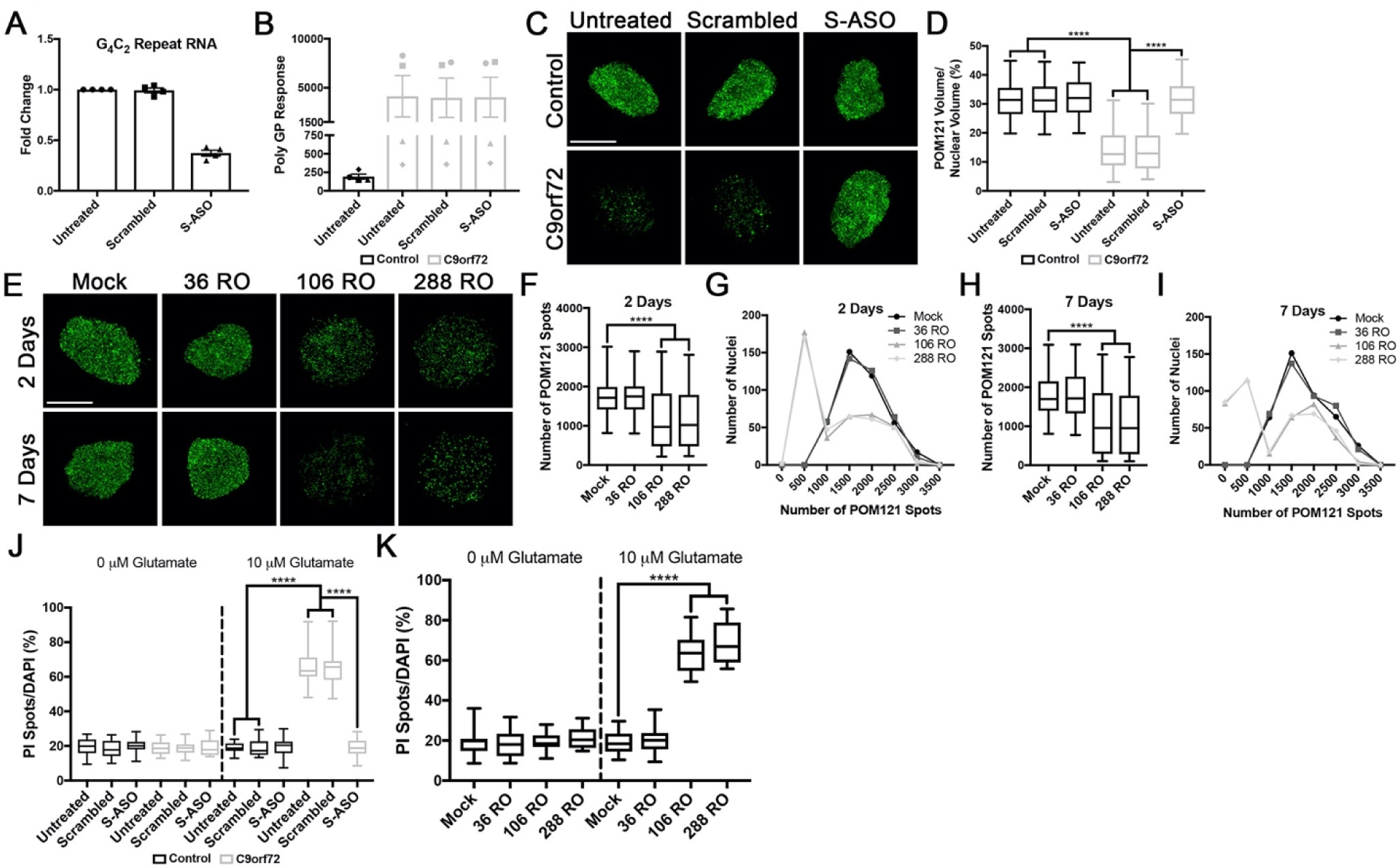
Expression of pathologic G_4_C_2_ repeat RNA initiated the nuclear loss of POM121. (**A-B**) qRT-PCR for G_4_C_2_ repeat RNA (**A**) and MSD Elisa for Poly(GP) DPR levels (**B**) in *C9orf72* iPSNs following 5 days of treatment with G_4_C_2_ targeting ASO. (**C**) SIM imaging for POM121 in nuclei isolated from control and *C9orf72* iPSNs. Genotype as indicated on left, treatment as indicated on top. (**D**) Quantification of percent total nuclear volume occupied by POM121 following 5 days of treatment with G_4_C_2_ targeting ASO. N = 4 control and 4 *C9orf72* IPSC lines, 50 NeuN+ nuclei per line/treatment. Two-way ANOVA with Tukey’s multiple comparison test was used to calculate statistical significance. **** p < 0.0001. (**E**) Maximum intensity projections from SIM imaging of POM121 in nuclei isolated from wildtype iPSNs overexpressing 36, 106, or 288 G_4_C_2_ RNA repeats. Time point as indicated on left, overexpression as indicated on top. (**F-I**) Quantification and histogram distributions of POM121 spots 2 days (**F-G**) and 7 days (**H-I**) after overexpression of G_4_C_2_ repeat RNA. N = 4 wildtype iPSC lines, 100 NeuN+ nuclei per line/overexpression. One-way ANOVA with Tukey’s multiple comparison test was used to calculate statistical significance. **** p < 0.0001. (**J**) Quantification of percent cell death following exposure to glutamate in iPSNs treated with ASOs for 5 days. N = 4 control and 4 *C9orf72* iPSC lines, 10 frames per well. Two-way ANOVA with Tukey’s multiple comparison test was used to calculate statistical significance. **** p < 0.0001. (**K**) Quantification of percent cell death following exposure to glutamate in wildtype iPSNs overexpressing G_4_C_2_ repeat RNA. N = 4 control iPSC lines, 10 frames per well. Two-way ANOVA with Tukey’s multiple comparison test was used to calculate statistical significance. **** p < 0.0001. Scale bar = 5 μm.

As an additional assessment to confirm the effects of G_4_C_2_ repeat RNA on the nuclear localization and expression of POM121, we transfected wildtype iPSNs with stop codon optimized constructs that produce G_4_C_2_ repeat RNA and generate pathologic RNA foci, but not DPRs, in immortalized cell lines (Mizielinska et al., 2014) and iPSNs (**Fig. S14)**. Following nuclei isolation and SIM, we found that overexpression of 106 or 288, but not 36, G_4_C_2_ RNA repeats in wildtype iPSNs resulted in a reduction in the nuclear localization and expression of POM121 after 2 and 7 days of G_4_C_2_ repeat RNA expression (**Fig. 6E-I**). Importantly, 7 day overexpression of CGG and CTG repeat RNAs (which cause Fragile X and DM1 muscular dystrophy, respectfully) in iPSNs did not induce the loss of POM121 from the nucleus (**Fig. S15**). Together, these data suggest that reduced nuclear expression and localization of POM121 in spinal neurons is a specific effect of pathological G_4_C_2_ repeat RNA. To assess the contribution of G_4_C_2_ repeat RNA itself to cell death, we performed glutamate toxicity assays. While treatment of *C9orf72* iPSNs with sense targeting ASOs for 5 days mitigated susceptibility to glutamate induced excitotoxicity (**Fig. 6J, Fig. S16A**), overexpression of 106 or 288 RNA repeats in wildtype iPSNs significantly increased susceptibility to glutamate induced excitotoxicity (**Fig. 6K, Fig. S16B**), mirroring the effects on POM121. Together, these data suggest that via direct or indirect mechanisms, G_4_C_2_ repeat RNA, but not DPRs or loss of C9ORF72 protein, initiates a pathological cascade impacting nuclear localization and expression of POM121 and subsequent cellular toxicity in human neurons.

## Discussion

Deficiencies in NCT have arisen as a prominent pathological mechanism underlying multiple neurodegenerative diseases including ALS, FTD, HD, and AD (Eftekharzadeh et al., 2018; Freibaum et al., 2015; Grima et al., 2017; Hutten and Dormann, 2019; Jovicic et al., 2015; Zhang et al., 2015). Although multiple studies have reported the nuclear or cytoplasmic mislocalization and accumulation of some NPC associated proteins in various model systems (Chew et al., 2019; Eftekharzadeh et al., 2018; Hutten and Dormann, 2019; Zhang et al., 2015; Zhang et al., 2018b; Zhang et al., 2016; Zhang et al., 2019b), few studies have linked these accumulations to alterations within the nucleus, nuclear envelope, and NPC itself. Furthermore, as many studies have focused on a handful of individual nucleoporins, a comprehensive evaluation of the expression and localization of individual Nups within the nucleus and NPCs is lacking. To evaluate the nuclear distribution, expression, and localization of the majority of individual human Nups, we harnessed the power and resolution of SIM. Here we studied >1,000,000 nuclei and comprehensively assessed actual nuclear repertoire of NPC components in nuclei from iPSNs as well as in postmortem human nervous system tissue from patients harboring the *C9orf72* HRE. We found that 8 of the 23 Nups analyzed are reliably and reproducibly lost from human neuronal nuclei in iPSNs (**Fig. 1**) and postmortem human motor cortex (**Fig. 2**) and spinal cord (**Fig. S6**). Notably, the stable NPC structure is formed early in cellular development and its individual Nup components are extremely long lived with some having half-lives measured in years (Savas et al., 2012; Toyama et al., 2013). The evaluation of Nups at multiple time points in the lifespan of our iPSNs reveals that initial nuclear localization and expression of individual Nups is not disrupted by the *C9orf72* HRE, but there is an age-related loss of specific Nups from the nucleus (**Fig. 1**). Although expression profiles and synaptic physiology of iPSNs more closely resemble those of young and immature fetal neurons as opposed to mature adult neurons (Ho et al., 2016), the age dependence (day 18 to day 32) of our observed alterations in specific Nups highlights the ability of our iPSN system to capture the maturation of early pathogenic cascades in *C9orf72* mediated neurodegeneration. Importantly, our iPSNs are not dying in culture without the addition of exogenous stressors and therefore our observed alterations in NPC components are not merely reflective of cell death cascades. Interestingly, although all 3 Nups of the transmembrane ring (GP210, NDC1, POM121) show decreased nuclear localization and expression, the remainder of the disrupted Nups span across multiple domains of the NPC including the nuclear basket (Nup50, TPR), central channel (Nup98), and outer ring (Nup107, Nup133). As these Nups collectively are involved in both maintaining the structural integrity and the function of the NPC, as well as NPC independent nuclear functions (Lin and Hoelz, 2019; Raices and D’Angelo, 2012), this suggests that *C9orf72* mediated alterations in individual NPC components themselves are likely to impact multiple cellular processes likely including both NCT and gene expression.

Although arranged into a highly organized octet structure, the NPC itself can be comprised of multiple copies (8, 16, 32 etc.) of each individual Nup and these numbers may vary depending on each Nup (Hampoelz et al., 2019; Kim et al., 2018; Lin and Hoelz, 2019). While SIM provides for increased resolution in detecting individual spots within isolated nuclei, it remains an open question as to whether this represents a singular or multiple Nup molecules within an NPC for each spot detected. Interestingly, there have been reports that the composition of each NPC within a single nucleus in a given cell can vary as individual nucleoporins are replaced and also varies amongst cell types (Kinoshita et al., 2012; Rajoo et al., 2018; Toyama et al., 2018). Given that little is known about the human neuronal nuclear pore complex, NPC heterogeneity within individual nuclei which may explain differences in Nup staining patterns. Additionally, it has been shown that the stoichiometry of the Nups within the NPC varies amongst different cell types (Ori et al., 2013; Rajoo et al., 2018). Future experiments will be needed to determine the exact stoichiometry of Nups within human neurons and individual NPCs. Nevertheless, these caveats do not undermine the consistent observation that a subset of transmembrane nucleoporins are reproducibly lost from human *C9orf72* neuronal nuclei *in vitro* and *in vivo*. Perhaps more importantly, the combinatorial loss of 8 specific Nups in the pathophysiology of *C9orf72* ALS directly impacts functional nuclear transport and downstream cellular physiology. Moreover, simply restoring nuclear envelope associated POM121, but not a soluble variant of POM121 not associated with the NPC, in neurons restores the entire nuclear composition of NPC components which together restore normal functional transport, and abrogates downstream stressor induced cytotoxicity.

Using the 414 antibody that recognizes 4 FG repeat containing Nups (Nup62, Nup153, Nup214, Nup358) and SIM, we show that the total number and distribution of nuclear pores is unchanged in *C9orf72* iPSNs (**Fig. S4A-C**) at the level of super resolution light microscopy. Together with our data that the nuclear localization and expression of each of the 8 altered Nups is on average decreased by about 50% (**Fig. 1E**), it is possible that the overall NPC is structurally intact but lacking 50% of the Nup molecules. Indeed, together with our 414 staining and SIM microscopy, our SEM results indicate that the distribution and architecture of NPCs in the nuclear envelope of *C9orf72* iPSNs remain intact (**Extended Data Fig. 4D**) supporting the hypothesis that a variable number of individual Nup molecules may be missing in each NPC without causing the collapse of the whole structure albeit our observed alterations are clearly impacting functional NCT. Alternatively, we can not rule out the possibility that some Nup spots detected by SIM are not associated with the NPC. Specific Nups have been shown to localize within the nucleus for NPC independent functions in gene expression and cell division (Raices and D’Angelo, 2012). While human neurons are no longer dividing, it is possible that some Nup spots detected by SIM are not associated with the NPC. The methods to reliably detect loss of a selected Nup from an individual NPC in human iPSNs remain unclear at this time.

Unlike previous studies based on overexpression of the *C9orf72* HRE or individual DPR species (Chew et al., 2019; Zhang et al., 2018b; Zhang et al., 2016; Zhang et al., 2019b), we do not observe mislocalization or aggregation of POM121 or other Nups (data not shown) in iPSNs (**Fig. S3C**) or human motor cortex (**Fig. S7**) harboring *endogenous levels* of *C9orf72* repeats and DPRs. In addition, Nup mRNA levels are unchanged in *C9orf72* iPSNs (**Fig. S3G**). Therefore, it is possible that as individual Nup molecules are lost from the nucleus, they are targeted for degradation in human neurons via the proteasome or autophagy. Future studies will be needed to uncover the specific pathways contributing to Nup loss and possible nuclear or cytoplasmic degradation in human neurons. Furthermore, it is equally possible that altered translation of Nups can compound the effects of their initial loss from the nucleus by preventing their nuclear replenishment.

The coordinated assembly of the NPC is well defined during mitosis and interphase in dividing cells and yeast (Lin and Hoelz, 2019; Otsuka and Ellenberg, 2018). Although the overall organization and structure of the NPC is fairly consistent amongst cell types (Lin and Hoelz, 2019), it is unknown whether functional relationships amongst specific Nups are conserved. Moreover, it is unclear whether disruptions in these associations may contribute to disease pathogenesis in neurons where many NPC components are extremely long lived and infrequently exchanged (Toyama et al., 2013). To begin to tackle this question in human neurons, we overexpressed candidate Nups from multiple domains of the NPC. Consistent with previous reports indicating that POM121 insertion into the nuclear envelope is a critical initiating event in NPC assembly during interphase (Antonin et al., 2005; Doucet et al., 2010; Funakoshi et al., 2011; Talamas and Hetzer, 2011), re-introduction of a single transmembrane Nup POM121 completely re-established the entire nuclear composition of individual Nups (**Fig 3A-T**). Interestingly, Nup133 overexpression only restored the nuclear localization and expression of non-transmembrane Nups (**Fig. 3A-T**). This is consistent with prior literature showing that Nup133 targets the Nup107-Nup160 Y complex to areas of inner and outer nuclear membrane fusion following POM121 insertion into the nuclear envelope (Doucet et al., 2010). Together, these data suggest that POM121 is critical for mediating the nuclear composition of individual Nups in human neurons.

The NPC is a barrier between the nucleus and cytoplasm to critically control NCT and cellular function (Lusk and King, 2017; Raices and D’Angelo, 2012). Thus, disruptions in the composition of Nups within the nucleus are likely to have detrimental effects on essential cellular processes including but not limited to NCT. Here we provide evidence that overexpression of POM121 re-establishes the nuclear expression and localization of specific Nups (**Fig. 3A-T**). The combined restoration of the nuclear expression and localization of affected Nups mitigates deficits in the subcellular distribution of Ran GTPase (**Fig. 3U-V**), functional active NCT (**Fig. S11**), and impacts downstream cellular toxicity in *C9orf72* iPSNs (**Fig. 3W**). In contrast, overexpression of Nup98, a Nup heavily involved in NCT itself through interactions between nuclear transport receptors and its FG repeat domain (Beck and Hurt, 2017; Chatel and Fahrenkrog, 2012; Raices and D’Angelo, 2012), has no effect on the localization of Ran (**Fig. 3U-V**). Together, this suggests that the complete composition of Nups within human neuronal nuclei is critical for maintenance of cellular function in human neurons.

To further probe the relationships between specific Nups in human neurons and their potential contribution to *C9orf72* mediated alterations in Nup composition and NPC function, we harnessed the power of Trim21 to rapidly degrade endogenous proteins (Clift et al., 2017).

Trim21 mediated degradation of POM121 in wildtype iPSNs recapitulates *C9orf72* mediated alterations in individual Nups (**Fig. 4A-P**). Together with our overexpression data, this suggests that loss of the transmembrane Nup POM121 is the initiating event in specific Nup alterations in human *C9orf72* neurodegeneration. We note that loss of POM121 alters the human neuronal nuclear repertoire of Nups and it is likely the combinatorial effect of these changes that impacts the subcellular distribution of Ran GTPase. Among other possibilities, disruption of the Ran gradient may be the result of alterations to Nups directly involved in NCT and/or a result of potential disruptions in gene transcription that may ultimately impact NCT. Future studies are necessary to define the impact of individual Nup alterations on specific aspects of functional NCT including those involving karyopherins and cargo loading/unloading. Together, the combined loss of 8 specific Nups and mislocalization of Ran GTPase are likely to impact multiple cellular pathways, the combination of which renders human neurons susceptible to stressor mediated cell death.

Intriguingly, POM121 overexpression or loss only affects those Nups that are disrupted in *C9orf72* iPSNs or human motor cortex suggesting that this specific subset of Nups (Nup50, TPR, GP210, NDC1, POM121, Nup107, Nup133) are physically, functionally, and/or mechanistically linked in human neurons. This is supported by studies in dividing cells suggesting functional and mechanistic links between a subset of these Nups during NPC assembly (Antonin et al., 2005; Beck and Hurt, 2017; Doucet et al., 2010; Lin and Hoelz, 2019; Mitchell et al., 2010; Otsuka and Ellenberg, 2018; Souquet et al., 2018).

Interestingly, the nuclear localization and expression of Nup98 is undisturbed following knockdown of either POM121 or Nup133. Moreover, nuclear localization and expression of Nup98 in partially restored upon overexpression of non-NPC associated sPOM121 (**Fig. S9A,D**) consistent with previous reports that sPOM121 localizes with nucleoplasmic Nup98 (Franks et al., 2016) where Nup98 plays a role as a transcriptional regulator (Kalverda et al., 2010; Liang et al., 2013; Raices and D’Angelo, 2017). Together, these data indicate that loss of Nup98 in *C9orf72* iPSN nuclei may be the result of a molecular mechanism independent of NPC associated POM121. Intriguingly, stress granule assembly has recently been shown to contribute to defects in NCT through the sequestration of Nups in dividing cells (Zhang et al., 2018a). Given that Nup98 is a highly mobile Nup which functions in the transport of molecules through the NPC (Griffis et al., 2002; Griffis et al., 2003), it is possible that its nuclear loss observed in *C9orf72* ALS is at least in part the result of incorporation into cytoplasmic stress granules.

It has previously been shown that Nups can modify *C9orf72* toxicity in *Drosophila* (Freibaum et al., 2015) and has been proposed that pathologic DPRs produced by the *C9orf72* HRE can interact with Nups to disrupt NCT (Lee et al., 2016; Shi et al., 2017). Here we found that in human neurons with endogenous expression of the *C9orf72* HRE, neither loss of C9ORF72 protein nor DPRs affect nuclear localization and expression of POM121 in human neurons (**Fig. 5A-E**). Recent studies have suggested that a reduction in DPR levels can mitigate stressor induced cell death and restore functional NCT in iPSNs (Cheng et al., 2019). However, this study did not evaluate the effects of DDX3X manipulation on NPC components themselves but was limited to the distribution of fluorescent shuttle proteins. Furthermore, we note that the mitigation of cell death and restoration of NCT was observed following overexpression of DDX3X, an RNA helicase involved in multiple aspects of RNA metabolism and export (Linder and Jankowsky, 2011; Tarn and Chang, 2009). As a result, it is possible that the protective effects of DDX3X overexpression are a result of altered RNA metabolism and not solely DPR expression. Although overexpression of DPRs does not negatively impact POM121, the possibility remains that DPRs may directly impact the localization of other NPC components (e.g. Nup98, NPC associated proteins). Furthermore, it is possible that DPRs may directly contribute to Nup independent alterations in functional NCT potentially via pathologic associations with karyopherins which have previously been identified as strong DPR interactors (Lee et al., 2016). Indeed, recent work from our lab shows that arginine rich DPRs Poly(GR) and Poly(PR) can directly interfere with importin beta function in non-neuronal systems (Hayes et al., 2019). As a result, DPRs dependent effects on the NCT machinery itself may further compound alterations in NCT that arise as a result of compromised nuclear expression and localization of Nups. Together, this could suggest a two-hit model for disruptions in NCT in *C9orf72* ALS/FTD. While DPRs themselves do not affect the nuclear expression and localization of POM121, we provide multiple lines of evidence suggesting that expression of G_4_C_2_ repeat RNA initiates the deficit in nuclear Nup composition and its downstream consequences. First, following the initial loss of POM121, 5 day exposure to a G_4_C_2_ targeting ASO greatly reduced G_4_C_2_ RNA but not Poly(GP) DPR levels and completely restored the nuclear localization and expression of POM121 (**Fig. 6A-D**) suggesting that G_4_C_2_ repeat RNA initiates a pathological cascade impacting POM121. Indeed, using stop codon optimized constructs which produce pathologic G_4_C_2_ RNA but not DPRs (Mizielinska et al., 2014), we show that overexpression of 106 or 288 G_4_C_2_ RNA not only results in reduced nuclear localization and expression of POM121 (**Fig. 6E-I**), but also increases susceptibility to glutamate induced excitotoxicity in these wildtype iPSNs (**Fig. 6K**). Thus, overexpressing just G_4_C_2_ repeat RNA in wildtype neurons, recapitulates our observations from patient iPSNs harboring endogenous levels of the *C9orf72* HRE. Notably, these results differ from previous studies in fly models overexpressing only G_4_C_2_ repeat RNA where no overt toxicity or degeneration was observed (Mizielinska et al., 2014). In the context of our current study in *human* spinal neurons, we note that flies do not have a gene encoding POM121 and thus may not be susceptible to this RNA meditated pathologic event.

Although, the loss of nuclear envelope associated POM121 is specific to the expression of G_4_C_2_ repeat RNA in spinal neurons (**Fig. 6, Fig. S15**), it still remains possible that CGG and/or CTG repeat RNAs can elicit similar deficits when expressed in their disease specific neuronal subtype. Together with our unpublished observations that overexpression of G_4_C_2_ repeat RNA in human astrocytes and HEK293T cells has no affect the nuclear localization or expression of POM121, our data support a role for the cell-type specific nature of G_4_C_2_ repeat RNA in the initiation of pathogenic events in human neurons. Collectively, these data suggest that the G_4_C_2_ repeat RNA itself mediates early disruptions in individual Nups. Future experiments are necessary to determine if this is the result of direct interactions between POM121 and G_4_C_2_ or G_2_C_4_ RNA as was shown to be the case for the NPC associated protein RanGAP1 (Zhang et al., 2015) or via indirect interacts via intermediate proteins and/or RNAs which may in turn affect the nuclear localization and expression of POM121. Our previous work did not identify POM121 as a direct interactor of G_4_C_2_ repeat RNA (Zhang et al., 2015), suggesting that *C9orf72* HRE mediated effects on the nuclear localization and expression of POM121 could be due to indirect interactions between G_4_C_2_ repeat RNA and POM121.

Together, our data support a role for G_4_C_2_ repeat RNA in early and initial pathogenic disruptions in ALS and additionally identify a function for POM121 in the maintenance of Nup composition within the nucleus in normal human neurons. Furthermore, our data highlight the importance of POM121 in the initiating events leading to overall disruptions in nuclear Nup composition, NPC function, and downstream impaired cellular survival in the pathogenesis of *C9orf72* ALS/FTD.

## Acknowledgements

We thank the ALS patients and their families for essential contributions to this research and the Target ALS Human Postmortem Tissue Core for providing postmortem human tissue. We also thank Martin Hetzer, Adrian Isaacs, Davide Trotti, and Maurice Swanson for kindly gifting POM121, G_4_C_2_ RNA Only, DPR, and (CTG)_202_ plasmids respectively. Expert technical assistance for iPSC maintenance was provided by Xiaopei Tang and Weibo Zhou. This work was supported by the ALSA Milton Safenowitz Postdoctoral Fellowship (ANC), along with funding from NIH-NINDS, The Robert Packard Center for ALS Research Answer ALS Program, ALS Finding a Cure, ALS Association, Muscular Dystrophy Association, and the Chan Zuckerberg Initiative.

## Author Contributions

Conceived and designed the experiments: ANC and JDR. Performed the experiments: ANC, BLZ, LH, BF, and HT. Analyzed the data: ANC, YZ, KB, HT, DS, and JDR. Contributed reagents and materials: ANC, BLZ, LH, FR, DS, CNS, AH, and JDR. Wrote the manuscript: ANC and JDR with input from co-authors.

## Experimental Procedures

### iPSC Neuron Differentiation

Peripheral blood mononuclear cell (PBMC)-derived iPSC lines from C9orf72-ALS patients and non-neurological controls were obtained from the Cedars-Sinai Answer ALS repository (see **Table S1** for demographics). Fibroblast derived isogenic iPSC lines were a kind gift from Kevin Talbot (Ababneh et al., 2019). The C9orf72 null line was a gift from Justin Ichida. iPSCs were maintained according to standard Cedars Sinai protocols and differentiated into spinal neurons according to the direct induced motor neurons (diMNs) protocol (**Fig. S1A**), which generates a mixed population consisting of 20-30% islet-1 positive motor neurons. Briefly, iPSC colonies were maintained on Matrigel coated 10 cm dishes for three weeks before passaging for differentiation. Once iPSC colonies reached 30-40% confluence (about 4-5 days after passaging), stage 1 media consisting of 47.5% IMDM (Gibco), 47.5% F12 (Gibco), 1% NEAA (Gibco), 1% Pen/Strep (Gibco), 2% B27 (Gibco), 1% N2 (Gibco), 0.2 μM LDN193189 (Stemgent), 10 μM SB431542 (StemCell Technologies), and 3 μM CHIR99021 (Sigma Aldrich) was added and exchanged daily until Day 6. On Day 6 of differentiation, cells were incubated in StemPro Accutase (Gibco) for 5 minutes at 37°C. Cells were collected from plates and centrifuged at 500 x g for 1.5 minutes. Cells were plated at 1 x 10^6 cells per well of a 6 well plate or 5 x 10^6 cells per T25 flask in stage 2 media consisting of 47.5% IMDM (Gibco), 47.5% F12 (Gibco), 1% NEAA (Gibco), 1% Pen/Strep (Gibco), 2% B27 (Gibco), 1% N2 (Gibco), 0.2 μM LDN193189 (Stemgent), 10 μM SB431542 (StemCell Technologies), 3 μM CHIR99021 (Sigma Aldrich), 0.1 μM all-trans RA (Sigma Aldrich), and 1 μM SAG (Cayman Chemicals). Media was exchanged daily until day 12. For the majority of experiments, on day 12 of differentiation, cells were switched to stage 3 media consisting of 47.5% IMDM (Gibco), 47.5% F12 (Gibco), 1% NEAA (Gibco), 1% Pen/Strep (Gibco), 2% B27 (Gibco), 1% N2 (Gibco), 0.1 μM Compound E (Millipore), 2.5 μM DAPT (Sigma Aldrich), 0.1 μM db-cAMP (Millipore), 0.5 μM all-trans RA (Sigma Aldrich), 0.1 μM SAG (Cayman Chemicals), 200 ng/mL Ascorbic Acid (Sigma Aldrich), 10 ng/mL BDNF (PeproTech), 10 ng/mL GDNF (PeproTech). For all whole cell imaging (TDP-43, Ran), cells were trypsinized and plated in 24 well optical bottom plates (Cellvis) at a density of 250,000 cells per well. Stage 3 media was exchanged every 3 days for the duration of the experiment. All cells were maintained at 37°C with 5% CO_2_ until day 18, 32, or 46 of differentiation as indicated by each experiment. iPSCs and iPSNs routinely tested negative for mycoplasma.

**Table S1:** Demographic Information for iPSC Lines

| **iPSC Line Name** | **Source** | **Clinical Diagnosis** | **Age at Time of Collection** | **Gender** | **Origin** |
| --- | --- | --- | --- | --- | --- |
| EDi036-A | Cedars-Sinai | Non-neurologic control | 79 | Female | PBMC |
| EDi037-A | Cedars-Sinai | Non-neurologic control | 79 | Male | PBMC |
| EDi029-A | Cedars-Sinai | Non-neurologic control | 80 | Male | PBMC |
| EDi034-A | Cedars-Sinai | Non-neurologic control | 79 | Female | PBMC |
| EDi022-A | Cedars-Sinai | Non-neurologic control | 79 | Male | PBMC |
| EDi044-A | Cedars-Sinai | Non-neurologic control | 80 | Female | PBMC |
| CS1ATZ | Cedars-Sinai | Non-neurologic control | 60 | Male | PBMC |
| CS8PAA | Cedars-Sinai | Non-neurologic control | 58 | Female | PBMC |
| EDi043-A | Cedars-Sinai | Non-neurologic control | 80 | Male | PBMC |
| CS0201 | Cedars-Sinai | Non-neurologic control | 56 | Female | PBMC |
| CS0BUU | Cedars-Sinai | C9orf72 | 63 | Female | PBMC |
| CS7VCZ | Cedars-Sinai | C9orf72 | 64 | Male | PBMC |
| CS0LPK | Cedars-Sinai | C9orf72 | 67 | Male | PBMC |
| CS6ZLD | Cedars-Sinai | C9orf72 |  | Female | PBMC |
| CS8KT3 | Cedars-Sinai | C9orf72 | 60 | Male | PBMC |
| CS2YNL | Cedars-Sinai | C9orf72 | 60 | Male | PBMC |
| CS0NKC | Cedars-Sinai | C9orf72 | 52 | Female | PBMC |
| CS6CLW | Cedars-Sinai | C9orf72 |  | Male | PBMC |
| 59-1 | K. Talbot | Isogenic Correction of OXC9-02 | 62 | Female | Fibroblast |
| OXC9-02-02 | K. Talbot | C9orf72 | 62 | Female | Fibroblast |
| 3231 | J. Ichida | Non-neurologic control (parent line for Null) | 56 | Male | PBMC |
| 3-11 | J. Ichida | C9orf72 Null | 56 | Male | PBMC |

### ASO Treatment of iPSC Derived Neurons

Scrambled ASO (676630): CCTATAGGACTATCCAGGAA and G_4_C_2_ ASO (619251): CAGGCTGCGGTTGTTTCCCT were provided by Ionis Pharmaceuticals. For 5 day ASO treatment, 5 μM scrambled or sense targeting ASO was added to the culture media on day 30. Media was exchanged and ASO replaced on day 32 and experiments were conducted on day 35 of differentiation.

### qRT-PCR

Samples were harvested in 1x DPBS with calcium and magnesium, then spun down in a microcentrifuge and the DPBS was aspirated. RLT Buffer (500 μL) was added to samples and RNA was isolated with an RNeasy kit (QIAGEN). RNA concentrations were determined using a NanoDrop 1000 spectrophotometer (Thermo Fisher Scientific). 1 μg RNA was used for cDNA synthesis using gene specific primers and a Superscript IV First-Strand cDNA Synthesis System (Thermo Fisher Scientific). qRT-PCR reactions were carried out using TaqMan Gene Expression Master Mix (TaqMan) and an Applied Biosystems Step One Plus Real Time PCR Machine (Applied Biosystems) using previously described primer/probe sets (see **Table S2** for sequences) (Lagier-Tourenne et al., 2013). Levels of *C9orf72* transcripts were normalized to GAPDH.

**Table S2:**
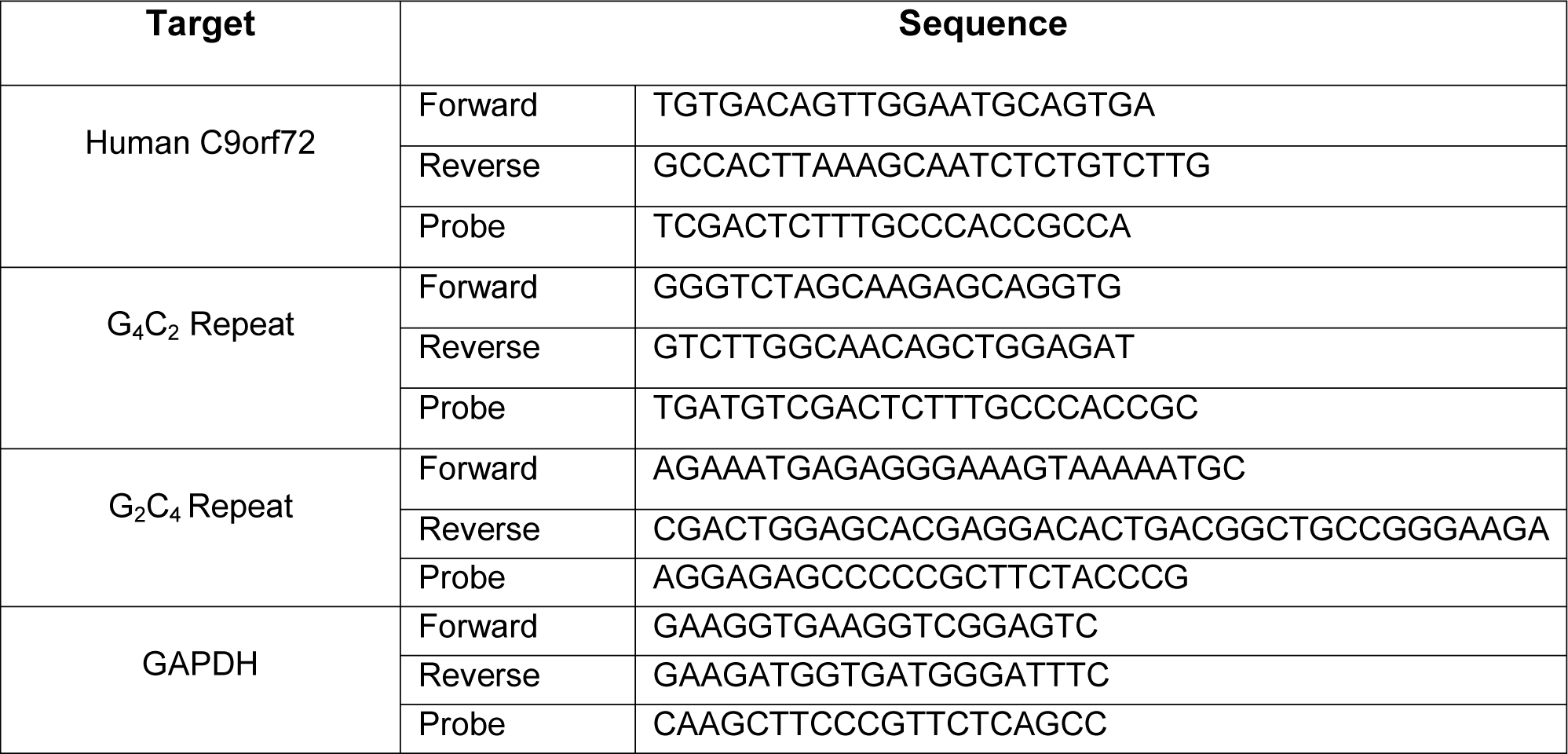
Primer and Probe Sequences for qRT-PCR

### Generation of poly-GP antibody

Two rabbits were immunized with the peptide antigen C-Ahx-(GP)_8_-amide (21^st^ Century Biochemicals). Specificity of antiserum versus pre-immune serum was verified by peptide dot blot and Western blot of HEK cells overexpressing GFP-tagged dipeptide repeat proteins (plasmids kindly provided by L. Petrucelli, Mayo) (Gendron et al., 2013). Antibodies were affinity purified before use.

### Meso Scale Discovery (MSD) ELISA

96-well small spot streptavidin coated plates were blocked overnight at 4° in 1% blocker A in PBS (all reagents from MSD). Following three washes in PBS-0.05% Tween (PBST), wells were coated with 0.35 µg/mL biotinylated rabbit anti-GP antibody for 1 hour at room temperature. After three PBST washes, cell lysates were added in duplicate, along with a standard curve of GP_8_ peptide. Following a 2-hour incubation at room temperature, wells were washed three times, and sulfo-tagged detection antibody added at 0.25 µg/mL for 1 hour. Following the final set of PBST washes, 150 µL read buffer was added and samples immediately imaged using MESO QuickPlex SQ 120. Specificity was verified using lysates of HEK cells overexpressing GFP-tagged dipeptide repeat proteins and linearity across range of interest assessed by serial dilution.

### Western Blots

Samples were harvested in 1x PBS, then spun down in a microcentrifuge and the PBS was aspirated. Samples were resuspended in RIPA buffer (Millipore Sigma) containing 1x protease inhibitor cocktail (Millipore Sigma). Samples were spun at 12,000g for 15 minutes to remove debris, and the supernatant was transferred to a new tube. 6x Laemmli buffer (12% SDS, 50% glycerol, 3% TrisHCl pH 7.0, 10% 2-mercaptoethanol in dH_2_O, bromophenol blue to color) was added to a final 1x concentration. Equal protein (by mass for quantifiable samples, by volume for unquantifiable samples) was loaded into 4-20% acrylamide gels and run until the dye front reached the bottom. Protein was transferred onto nitrocellulose membrane (Bio-Rad). Following transfer, blots were blocked for 1 hour in 5% milk in 1x TBS-Triton (0.1%). Blots were probed overnight (approximately 16 hours) at 4 degrees. Blots were washed four times in 1x TBST for 10 minutes each, probed with secondary antibody for 1 hour at room temperature, and washed another four times in 1x TBST for 10 minutes each. ECL substrate (Millipore Sigma, Thermo Fisher Scientific) was applied for 30 seconds, then images were taken using the GE Healthcare ImageQuant LAS 4000. Images were quantified using FIJI. For nuclei westerns, nucleoporins were normalized to total protein levels using the BLOT-FastStain Kit (G-Biosciences).

### Nanostring Gene Expression Analysis

RNA was quantified using a custom probe codeset (see **Table S3**) and was performed with 100-200 ng RNA using an nCounter Analysis System according to manufacturer’s protocol. RNA counts were normalized as previously described (Donnelly et al., 2013) using the nSolver Analysis Software (Nanostring) and a panel of control housekeeping genes (see **Table S3**).

**Table S3:**
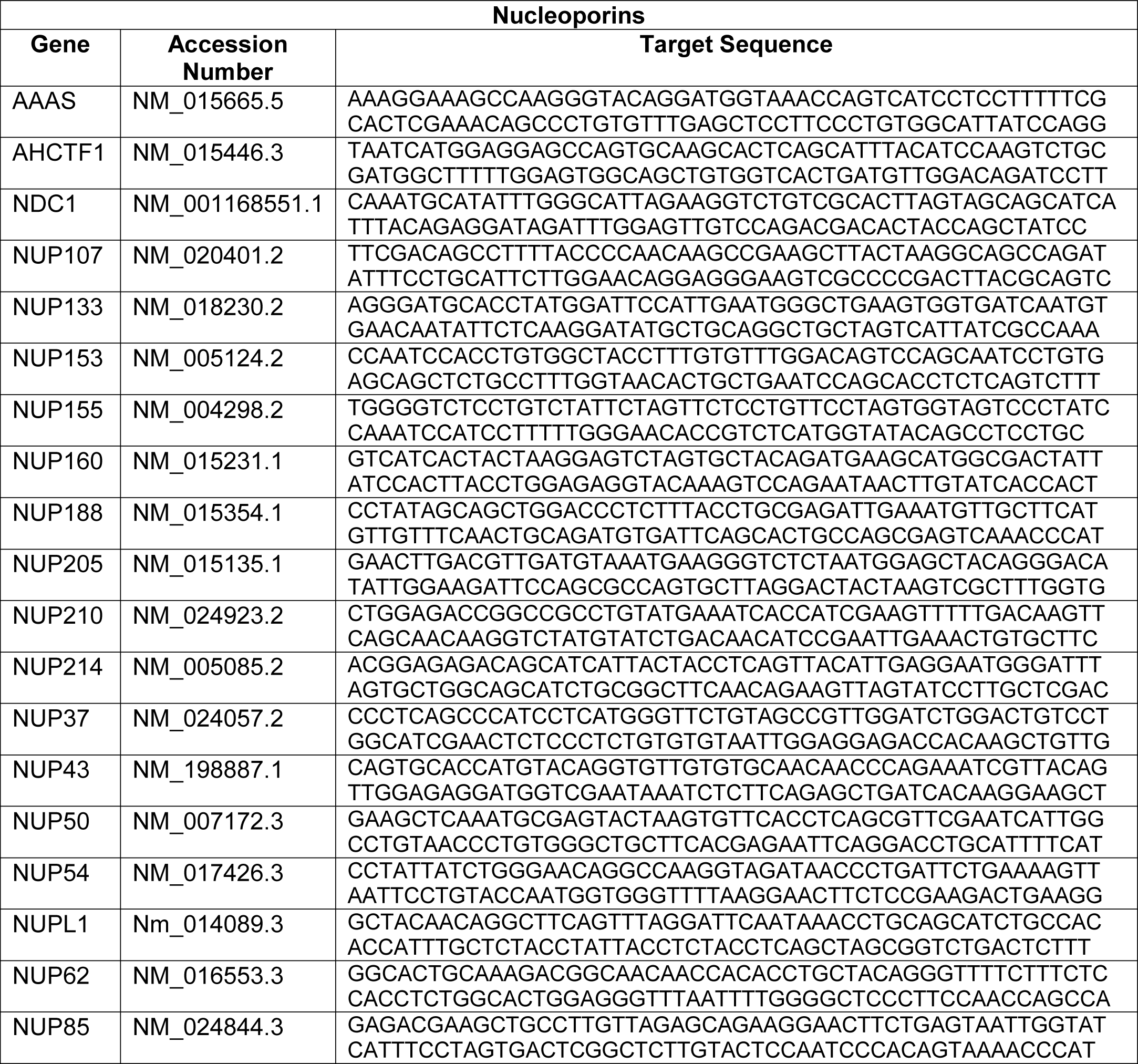
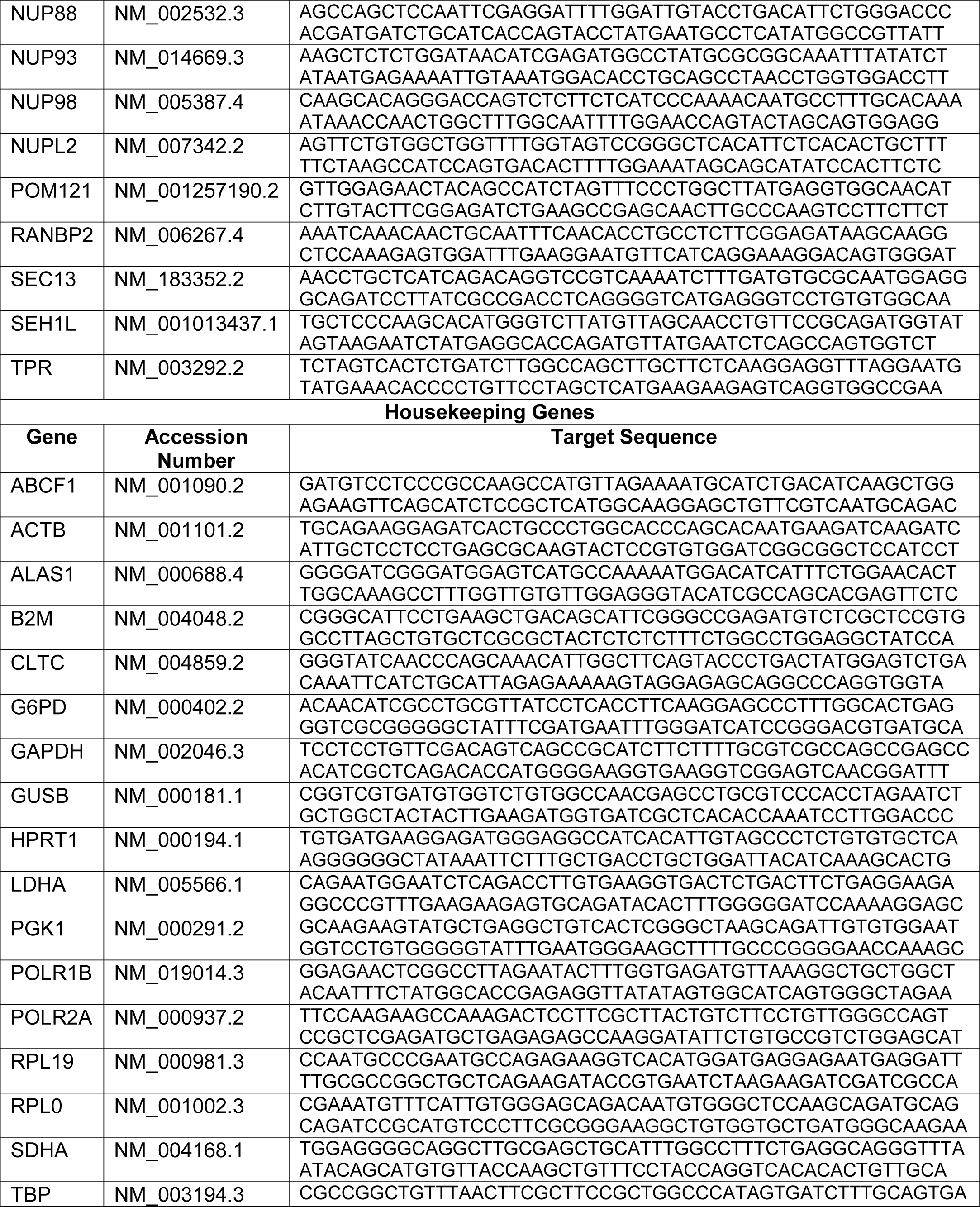
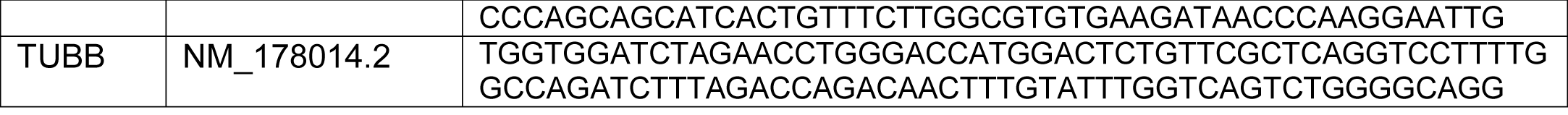
Custom Nanostring Codeset for Target Nucleoporins and Housekeeping Genes

### Nuclei Isolation and Super Resolution Structured Illumination Microscopy

All nuclei were isolated using Nuclei Pure Prep Nuclei Isolation Kit (Sigma Aldrich) using a sucrose gradient of 1.85 M. For preparation of iPSN lysates, media was aspirated from each well and cells were briefly rinsed with 1X PBS. iPSNs were scraped in lysis buffer using a cell scraper, transferred to a 50 mL conical tube, and vortexed. For preparation of postmortem human brain lysates, 75-100 mg of fresh frozen tissue was homogenized in lysis buffer using a dounce homogenizer. Sucrose gradients were assembled and centrifuged following the Nuclei Pure Prep Nuclei Isolation Kit protocol (Sigma Aldrich). Samples were centrifuged at 15,600 rpm and 4°C using a Swi32T swinging bucket rotor and Beckman ultracentrifuge (Beckman Coulter) for 45 minutes. Following isolation, nuclei were resuspended in 1 mL of Nuclei Storage Buffer and 25-100 μL was spun onto slides coated with 1 mg/mL collagen (Advanced Biomatrix) using a CytoSpin 4 Centrifuge (Thermo Fisher Scientific). Nuclei were fixed in 4% PFA for 5 minutes, washed with 1X PBS 3X 10 minutes, permeabilized with 1X PBST containing 0.1% Triton X-100 for 15 minutes, blocked in 10% normal goat serum diluted in 1X PBS for 1 hour, and incubated in primary antibody overnight at 4°C (See **Table S4** for antibody information). The next day, nuclei were washed with 1X PBS 3X 10 minutes, incubated in secondary antibody (See **Table S4** for antibody information) for 1 hour at room temperature, and washed in 1X PBS 3X 10 minutes. Nuclei were coverslipped using Prolong Gold Antifade Reagent (Invitrogen) and 18 mm x 18 mm 1.5 high tolerance coverslips (MatTek). NeuN or GFP positive nuclei were imaged by super resolution structured illumination microscopy (SIM) using a Zeiss ELYRA S1. Nucleoporin spots and volume quantification was automated using Imaris version 9.2.0 (Bitplane) and the 3D suite in FIJI as previously described (Eftekharzadeh et al., 2018). Briefly, nucleoporin spots were counted using automated spot detection in which a Bayesian classifier which takes into account features such as volume, average intensity, and contrast was applied to detect and segment individual spots. The total number of nucleoporin spots was determined using a 3D-rendering of super resolution SIM images where z-slices were taken through the entire nucleus. When individual nucleoporin spots could not be resolved due to limits of resolution of immunofluorescent SIM, the percent total nuclear volume occupied by the nucleoporin was calculated. X and Y axis length was measured in the center z-slice for each nucleus and Z axis length was estimated by multiplying z-slice thickness by the number of z slices imaged for each nucleus. These measurements were used to calculate total nuclear volume. For volume of the nucleoporin, automatic thresholding was applied and 3D suite in FIJI was used to determine the volume of the thresholded area for each nucleus. For a subset of images, automated 3D volume calculations were verified by measuring the area of the thresholded nucleoporin signal for each z-slice taken per nucleus. Z-slice thickness was then used to determine the volume of the nucleoporin for each individual z-slice. The resulting values were combined to yield total nucleoporin volume per nucleus. Nucleoporin volume was divided by total nuclear volume to yield a percentage of the total nuclear volume occupied by a given nucleoporin. In total across all experiments, over 1,000,000 nuclei were imaged and analyzed. Images presented are 3D maximum intensity projections generated in Zeiss Zen Black 2.3 SP1.

**Table S4:** Antibody Information

| Primary Antibodies |  |  |  |
| --- | --- | --- | --- |
| Antibody | Source | Catalog Number | Application and Concentration |
| Rabbit Anti-Nup50 | Abcam | ab137092 | IF: 1/250 |
| Rabbit Anti-Nup153 | Abcam | ab84872 | IF: 1/200 |
| Mouse Anti-TPR | Santa Cruz Biotechnology | sc-271565 | IF: 1/50 |
| Rabbit Anti-Nup54 | Abcam | ab220890 | IF: 1/250 |
| Mouse Anti-Nup58 (NUPL1) | Abcam | ab69954 | IF: 1/200 |
| Rat Anti-Nup62 | Millipore | MABE1043 | IF: 1/250 |
| Rat Anti-Nup98 | Abcam | ab50610 | IF: 1/250 |
| Rabbit Anti-GP210 | Abcam | ab15601 | IF: 1/200 |
| Rabbit Anti-NDC1 (TMEM48) | Thermo Fisher Scientific | PA5-67130 | IF: 1/200 |
| Rabbit Anti-NDC1 (TMEM48) | Novus Biologicals | NBP1-91603 | Western: 1/1000 |
| Rabbit Anti-POM121 (C Terminal) | Thermo Fisher Scientific | PA5-36498 | IF: 1/250 |
| Rabbit Anti-POM121 (N Terminal) | Novus Biologicals | NBP2-19890 | IF: 1/250<br>Western: 1/1000 |
| Mouse Anti-Nup93 | Santa Cruz Biotechnology | sc-374399 | IF: 1/50 |
| Rabbit Anti-Nup155 | Abcam | ab199528 | IF: 1/200 |
| Rabbit Anti-Nup188 | Thermo Fisher Scientific | PA5-63984 | IF: 1/250 |
| Mouse Anti-Nup205 | Santa Cruz Biotechnology | sc-377047 | IF: 1/50 |
| Rabbit Anti-ELYS (AHCTF1) | Thermo Fisher Scientific | PA5-56812 | IF: 1/200 |
| Rabbit Anti-Nup37 | Thermo Fisher Scientific | PA5-66007 | IF: 1/200 |
| Rabbit Anti-Nup43 | Abcam | ab69447 | IF: 1/200 |
| Mouse Anti-Nup107 | Thermo Fisher Scientific | MA110031 | IF: 1/200 |
| Mouse Anti-Nup133 | Santa Cruz Biotechnology | sc-376699 | IF: 1/50<br>Western: 1/1000 |
| Rabbit Anti-Nup160 | Abcam | ab74147 | IF: 1/250 |
| Rabbit Anti-Nup88 | Abcam | ab79785 | IF: 1/250 |
| Rabbit Anti-Nup214 | Bethyl Laboratories | A300-716A | IF: 1/200 |
| Mouse Anti-Nup358 (RANBP2) | Santa Cruz Biotechnology | sc-74518 | IF: 1/50 |
| Mouse Anti-414 | Abcam | ab50008 | IF: 1/250 |
| Chicken Anti-NeuN | Millipore | ABN91 | IF: 1/500 |
| Chicken Anti-GFP | Millipore | AB16901 | IF: 1/1000 |
| Rabbit Anti-TDP-43 | ProteinTech | 10782-2-AP | IF: 1/250 |
| Mouse Anti-TDP-43 | Abcam | ab104223 | IF: 1/250 |
| Guinea Pig Anti-Map2 | Synaptic Systems | 188004 | IF: 1/1000 |
| Mouse Anti-Ran | BD Biosciences | 610341 | IF: 1/200 |
| Mouse Anti-SMI32 | Covance | SMI-32P | IF: 1/1000 |
| Mouse Anti-Tuj1 | Sigma Aldrich | MAB1637 | IF: 1/1000-1/2000 |
| Rat Anti-NKX6.1 | DSHB | F55A12 | IF: 1/50 |
| Goat Anti-Islet1 | R&D Systems | AF1837 | IF: 1/100 |
| Rabbit Anti-C9ORF72 | ProteinTech | 22637-1-AP | Western: 1/1000 |
| Mouse Anti-GAPDH | Life Technologies | AM4300 | Western: 1/10,000 |
| Rabbit Anti-Alpha Tubulin | Cell Signaling | 2125 | Western: 1/1000 |
| **Secondary Antibodies** |  |  |  |
| Goat Anti-Mouse Alexa 488 | Invitrogen | A11029 | IF: 1/1000 |
| Goat Anti-Rabbit Alexa 488 | Invitrogen | A11034 | IF: 1/1000 |
| Goat Anti-Rat Alexa 488 | Invitrogen | A11006 | IF: 1/1000 |
| Goat Anti-Chicken Alexa 488 | Invitrogen | A11039 | IF: 1/1000 |
| Goat Anti-Mouse Alexa 568 | Invitrogen | A11031 | IF: 1/1000 |
| Goat Anti-Rabbit Alexa 568 | Invitrogen | A11036 | IF: 1/1000 |
| Goat Anti-Rat Alexa 568 | Invitrogen | A11077 | IF: 1/1000 |
| Goat Anti-Guinea Pig Alexa 568 | Invitrogen | A11075 | IF: 1/1000 |
| Goat Anti-Mouse Alexa 647 | Invitrogen | A21236 | IF: 1/1000 |
| Goat Anti-Rabbit Alexa 647 | Invitrogen | A21245 | IF: 1/1000 |
| Goat Anti-Rat Alexa 647 | Invitrogen | A21247 | IF: 1/1000 |
| Goat Anti-Chicken Alexa 647 | Invitrogen | A21449 | IF: 1/1000 |
| Goat Anti-Guinea Pig Alexa 647 | Invitrogen | A21450 | IF: 1/1000 |
| Donkey Anti-Mouse Alexa 488 | Invitrogen | A21202 | IF: 1/1000 |
| Donkey Anti-Rat Alexa 488 | Invitrogen | A21208 | IF: 1/1000 |
| Donkey Anti-Mouse Alexa 568 | Invitrogen | A10037 | IF: 1/1000 |
| Donkey Anti-Goat Alexa 568 | Invitrogen | A11057 | IF: 1/1000 |
| Donkey Anti-Rabbit IgG HRP | Thermo Fisher Scientific | 45-000-682 | Western: 1/5000 |
| Horse Anti-Mouse IgG HRP | Cell Signaling | 7076S | Western: 1/5000 |

### Immunofluorescent Staining and Confocal Imaging

On day 12 of differentiation, iPSNs were plated in 24 well optical bottom plates (Cellvis). iPSNs were fixed in 4% PFA for 15 minutes, washed with 1X PBS 3X 10 minutes, permeabilized with 1X PBST containing 0.3% Triton X-100 for 15 minutes, blocked in 10% normal goat serum diluted in 1X PBS for 1 hour, and incubated in primary antibody for 2 hours at room temperature (See **Table S4** for antibody information). iPSNs were then washed with 1X PBS 3X 10 minutes, incubated in secondary antibody (See **Table S4** for antibody information) for 1 hour at room temperature, washed in 1X PBS 2X 10 minutes, incubated with Hoescht diluted 1:1000 in 1X PBS for 10 minutes, and washed in 1X PBS 2X 10 minutes. iPSNs were mounted using Prolong Gold Antifade Reagent with DAPI. iPSNs were imaged using a Zeiss LSM 800 confocal microscope and nuclear intensity and N/C ratios were calculated with FIJI as previously described (Zhang et al., 2015). Images presented are maximum intensity projections generated in Zeiss Zen Blue 2.3.

### Human Tissue Immunofluorescence

Postmortem paraffin embedded motor cortex (See **Table S5** for demographic information) was rehydrated with xylene 3X 5 mins, 100% ethanol 2X 5 mins, 90% ethanol 5 mins, 70% ethanol, dH_2_O 3X 5 mins. Antigen retrieval was performed using Tissue-Tek antigen retrieval solution (IHC World) for 1 hour in a steamer. Slides were cooled for 10 mins, washed 3X 5 mins with dH_2_O followed by 2X 5 mins in 1X PBS. Tissue was permeabilized with 0.4% Triton X-100 in 1X PBS for 10 mins with gentle agitation on a shaker. Slides were washed 3X 5 mins with 1X PBS and blocked in DAKO protein-free serum block (DAKO) overnight in a humidified chamber at 4°C. Tissue was incubated in primary antibody diluted in DAKO antibody diluent reagent with background reducing components (DAKO) for 12 hours at room temperature with gentle agitation on a shaker followed by 2 days at 4°C (See **Table S4** for antibody information). Slides were washed 3X 5 mins in 1X PBS and incubated in secondary antibody diluted in DAKO antibody diluent reagent with background reducing components (DAKO) for 1 hour at room temperature with gentle agitation on a shaker (See **Table S4** for antibody information). Slides were washed 3X 5 mins in 1X PBS, rinsed briefly with autofluorescence eliminator reagent (Millipore), washed 5X 5 mins in 1X PBS, stained with Hoescht diluted 1:1000 in 1X PBS for 20 mins, and washed 3X 5 mins in 1X PBS. Slides were coverslipped using Prolong Gold Antifade Reagent with DAPI and nuclei from Map2 positive Layer V neurons were imaged with a Zeiss Axioimager Z2 fluorescent microscope equipped with an apotome2 module. POM121 nuclear intensity was quantified using FIJI. Images presented are maximum intensity projections generated in Zeiss Zen Blue 2.3.

**Table S5:** Demographic Information for Human Tissue

|  | **Clinical Diagnosis** | **Age of Death** | **Gender** |
| --- | --- | --- | --- |
| **Control-1** | Non-neurologic control |  | Female |
| **Control-2** | Non-neurologic control | 70 | Female |
| **Control-3** | Non-neurologic control | 70 | Male |
| **Control-4** | Non-neurologic control | 59 | Male |
| **Control-5** | Non-neurologic control | 71 | Female |
| **Control-6** | Non-neurologic control | 52 | Male |
| **Control-7** | Non-neurologic control | 72 | Male |
| **Control-8** | Non-neurologic control | 74 | Female |
| **C9orf72-1** | C9orf72 | 72 | Male |
| **C9orf72-2** | C9orf72 | 69 | Female |
| **C9orf72-3** | C9orf72 | 61 | Female |
| **C9orf72-4** | C9orf72 | 51 | Female |
| **C9orf72-5** | C9orf72 | 59 | Male |
| **C9orf72-6** | C9orf72 | 61 | Female |
| **C9orf72-7** | C9orf72 | 68 | Female |

### Plasmids and Nucleofection for Nup, G_4_C_2_ repeat RNA, and DPR Overexpression

iPSNs were transfected using the Lonza nucleofection system (program DC104) and a Lonza P3 Primary Cell 4D Nucleofector Kit (Lonza). POM121, Nup133, NDC1, GP210, and Nup98 plasmids were obtained from Origene. Additional POM121 and sPOM121 plasmids (Franks et al., 2016) were a kind gift from Martin Hetzer. GFP plasmids were obtained from Origene and Lonza. Poly(GA)_50_, Poly(GR)_50_, and Poly(PR)_50_ DPR plasmids (Wen et al., 2014) were a kind gift from Davide Trotti and (G_4_C_2_)_n_ RNA only plasmids (Mizielinska et al., 2014) were a kind gift from Adrian Isaacs. The (CGG)_99_ plasmid was obtained from Addgene and the (CTG)_202_ plasmid was a kind gift from Maurice Swanson. For Nup overexpression experiments, 5 x 10^6^ iPSNs were nucleofected with 4 μg of DNA on day 30 of differentiation. Expression was observed at day 31 and iPSNs were assayed at day 46 of differentiation. For DPR overexpression experiments, 5 x 10^6^ iPSNs were nucleofected with 4 μg or 1 μg of DNA on day 18 and assayed at day 20 or 25 of differentiation respectively. For RNA only experiments, 5 x 10^6^ iPSNs were nucleofected with 4 μg or 1 μg of DNA on day 18 and assayed at day 20 or 25 of differentiation respectively. For CGG and CTG overexpression experiments, 5 x 10^6^ iPSNs were nucleofected with 1 μg of DNA on day 18 and assayed at day 25 of differentiation.

### Knockdown of Nups by Trim Away

Trim21 mediated knockdown of Nups was modified from a previously described protocol (Clift et al., 2017). POM121 (Thermo Fisher Scientific), Nup133 (Santa Cruz Biotechnology), NDC1 (Novus Biologicals), and GAPDH (Cell Signaling) antibodies were dialyzed using Slide-A-Lyzer MINI Dialysis Device, 20K MWCO (Thermo Fisher Scientific). Following dialysis, antibodies were concentrated using Amicon Ultra-0.5 Centrifugal Filter Unit with Ultracel-100 membrane (Millipore). Antibody concentration was determined using a NanoDrop 1000 spectrophotometer (Thermo Fisher Scientific) as described (Clift et al., 2017). On day 18 of differentiation, 5 x 10^6^ iPSNs were nucleofected with 5 μg of antibody and 4 μg Trim21 GFP plasmid DNA (Origene) with Lonza P3 Primary Cell 4D Nucleofector Kit (Lonza) using program DC154. The next day, Trim21 GFP expression was observed and iPSNs were assayed on day 20 of differentiation.

### NLS-tdTomato-NES (S-tdTomato) Nucleocytoplasmic Transport Assay

On day 32 of differentiation, control and *C9orf72* iPSNs were nucleofected using the Lonza nucleofection system (program DC104) and a Lonza P3 Primary Cell 4D Nucleofector Kit (Lonza) as described above. One day later, on day 33 of differentiation, iPSNs were transduced with NLS-tdTomato-NES (S-tdTomato) lenti virus as previously described (Zhang et al., 2015). On days 34, 36, and 43 of differentiation (1, 3, and 10 or 2, 5, and 11 days post S-tdTomato transduction or POM121 OE respectively), iPSNs were fixed with 4% PFA for 15 mins, washed 3X 10 mins with 1X PBS and then incubated with Hoescht diluted 1:1000 in 1X PBS for 10 minutes, and washed in 1X PBS 2X 10 minutes. iPSNs were then immediately imaged using a Zeiss LSM 800 confocal microscope and the N/C ratio of S-tdTomato was calculated with FIJI as previously described (Zhang et al., 2015).

### Glutamate Toxicity

On day 12 of differentiation, iPSNs were plated in 24 well optical bottom plates (Cellvis) at a density of 250,000 neurons per well. Neurons were rinsed with 1X PBS and fed with fresh stage 3 media daily remove dead cells and debris prior to incubation with glutamate. For POM121 overexpression experiments, iPSNs were split, transfected as described above, and plated into optical bottom plates on day 30 of differentiation. For these experiments, iPSNs were washed/fed daily from day 32-46 before the toxicity experiments were performed. For 5 day ASO treatment experiments, iPSNs were treated with 5 μM ASO on day 30 and 32 during which time they were washed with 1X PBS. For G_4_C_2_ Repeat RNA only experiments, iPSNs were split and transfected with 1 μg of DNA on day 18 as described above. iPSNs were washed/fed daily from day 19-25 before the toxicity experiments were performed. On the day of the experiment (day 46 for POM121 OE, day 35 for 5 Day ASO, day 25 for G_4_C_2_ Repeat RNA only), iPSNs were washed with 1X PBS to remove any remaining debris and dead cells. Media was replaced with ACSF (Tocris) containing 0, 1, 10, or 100 μM glutamate (Sigma Aldrich). iPSNs were incubated at 37°C with 5% CO_2_ for 4 hours. After 3.5 hours, one drop of NucBlue Live ReadyProbes (Thermo Fisher Scientific) and 1 μM propidium iodide (Thermo Fisher Scientific) and returned to the incubator for 30 mins. iPSNs were imaged in an environmentally controlled chamber with a Zeiss LSM 800 confocal microscope. 10 images were taken and analyzed per well. PI and DAPI spots were counted using FIJI. For POM121 overexpression experiments, following live imaging, PI and NucBlue were washed out of neurons and iPSNs were subsequently immunostained for GFP and Map2 as described above and imaged with a Zeiss LSM 800 confocal microscope.

### Scanning Electron Microscopy

Previously published protocols (Fichtman et al., 2019; Fichtman et al., 2014) were modified to expose iPSN nuclei for high resolution imaging. Briefly, day 32 control and *C9orf72* iPSNs were frozen in growth medium supplemented with 10% DMSO, shipped on dry ice, and stored under liquid nitrogen until use. Thawed cells were pelleted, resuspended in mildly hypotonic buffer in the absence of detergents, and gently passed through a 21-gauge needle. The specimens were then centrifuged onto the surface of 5 x 5 mm^2^ silicon chips (Ted Pella), fixed, and further processed for electron microscopy as previously described (Fichtman et al., 2019; Fichtman et al., 2014), including controlled dehydration through a graded series of ethanol solutions and critical-point drying on a K850 apparatus (Quorum Technologies). The specimens were then coated with a ∼1 nm thick layer of iridium using a Q150T turbo-pumped sputter coater (Quorum Technologies) and imaged on a *Merlin* field emission scanning electron microscope (Zeiss) equipped with a secondary electron in-lens detector.

### Dextran Exclusion

On day 32 of differentiation, iPSNs were rinsed briefly with 1X PBS to remove remaining traces of media. iPSNs were permeabilized in permeabilization buffer (10 mM Tris pH 7.5, 10% OptiPrep, ultrapure water) containing 200 μg/mL digitonin on ice for 5 mins. iPSNs were then rinsed with ice cold transport buffer (4 mM HEPES-KOH pH 7.5, 22 mM KOAc, 0.4 mM Mg(OAc)_2_, 1 mM NaOAc, 0.1 mM EGTA, 50 mM sucrose) 2X 5 mins. Permeabilized iPSNs were incubated with 0.6 mg/mL fluorescent dextrans diluted in transport buffer for 25 mins at room temperature protected from light. During this time, one drop of NucBlue was added to each well. Following incubation with dextrans and NucBlue, iPSNs were imaged using a Zeiss LSM 800 confocal microscope and a 40X objective. Nuclear intensity of fluorescent dextrans was determined with FIJI.

### RNA FISH

Following one week of (G_4_C_2_)_n_ repeat RNA overexpression, iPSNs were rinsed briefly in 1X PBS and fixed in 4% paraformaldehyde in 1X PBS for 15 minutes at room temperature. iPSNs were then rinsed 3X 5 mins with 1X PBS, permeabilized with 1X PBS containing 0.3% Triton X-100 for 15 mins at room temperature, and subsequently washed 3X 5 mins with 1X PBS. iPSNs were then equilibrated in 1X SSC for 10 mins at room temperature. For RNase A treatment, 40 μg/mL RNase A in 1X SSC was added to corresponding wells and iPSNs were incubated at 37°C for 45 mins. RNase treated iPSNs were washed briefly 2X with 1X SSC and then equilibrated in 1X SSC for 10 mins at room temperature. iPSNs were then equilibrated in 50% formamide in 2X SSC for 10 mins at 60°C and then hybridized with hybridization buffer (40% formamide, 2 mg/mL heat shock inactivated BSA, 1 mM ribonucleoside-vanadyl complex, 10 mM NaPO_4_, 1X SSC) and probe mixture (10 μg salmon sperm DNA, 10 μg E.coli tRNA, 80% formamide, 0.2 μM custom LNA probe) for 1 hour at 60°C. Custom 5’ DIG labeled LNA probes (Qiagen) are as follows: G_4_C_2_ probe: CCCCGGCCCCGGCCCC lot number 302094012, scrambled probe: GTGTAACACGTCTATACGCCCA lot number 302495047. iPSNs were then washed 1X 20 mins with 50% formamide in 2X SSC at 65°C, 1X 15 mins with fresh 50% formamide in 2X SSC at 65°C, 1X 10 mins with 1X SSC in 40% formamide at 60°C, 3X briefly with 1X SSC at room temperature, 2X 5 mins with 1X SSC at room temperature, 1X 5 mins with TBS (50 mM Tris pH 7.5, 15 mM NaCl), 1X 5 mins with Tris Glycine (0.75% glycine, 0.2 M Tris pH 7.5), post-fixed with 3% paraformaldehyde in 1X PBS, and briefly rinsed 3X with 1X PBS. iPSNs were blocked with 1% normal goat serum and 5% heat shock inactivated BSA in TBS for 1 hour at room temperature and then incubated in 1:500 Mouse Anti-DIG (Jackson Immunoresearch) diluted in immunofluorescence buffer (2% heat shock inactivated BSA, TBS) overnight at 4°C. iPSNs were then washed 3X 5 mins with immunofluorescence buffer at room temperature and incubated with secondary antibody (1:500 Goat Anti-Mouse Alexa 568) diluted in immunofluorescence buffer for 30 mins at room temperature before undergoing a series of washes: 1X 5 mins TBS, 1X 5 mins Tris Glycine, 1X 5 mins PBS MgCl_2_ (2 mM MgCl_2_, 1X PBS), 1X 5 mins PBS, 1X 5 mins PBS with 1:1000 Hoescht, 1X 5 mins PBS. iPSNs were mounted with prolong gold antifade reagent with DAPI and imaged using a Zeiss LSM 800 confocal microscope and 63X objective. All solutions were made using RNase free water.

### Statistical Analysis

All data analysis was conducted with Imaris version 8.3.1 or FIJI as described above. All statistical analyses were performed using GraphPad Prism version 7 (GraphPad). Statistical analyses were performed with regards to individual iPSC lines or patients such that the average of all nuclei or cells evaluated per iPSC line or patient represents N = 1 with total N per experiment and group as indicated in figure legends. Student’s T-test, One-way ANOVA with Tukey’s multiple comparison test, or Two-way ANOVA with Tukey’s multiple comparison test was used as described in figure legends. * p < 0.05, ** p < 0.005, *** p < 0.0005, **** p < 0.0001. Box and whisker plots used to display the full spread of large data sets (>10 data points). Box encompasses the 25^th^ to 75^th^ percentile with the center bar identifying the mean. Whiskers represent min to max data points. For smaller data sets, bar graphs displaying individual data points shown where error bars represent +/- SEM.

---

**Figure S1.**
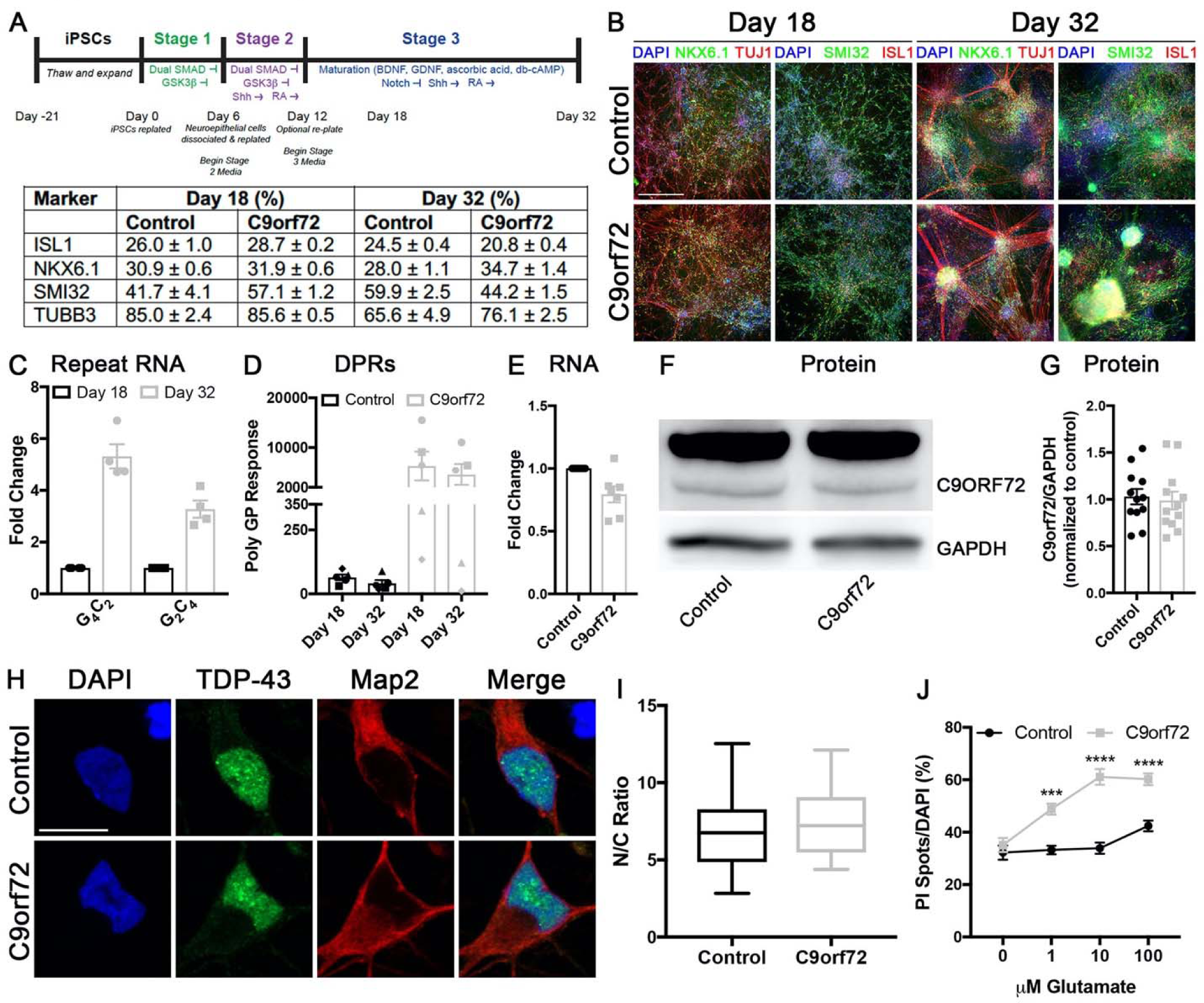
Related to Figure 1: *C9orf72* pathology is present in an accelerated iPSC derived neuron differentiation protocol. (**A**) Schematic of spinal neuron differentiation protocol and table of cell type marker quantification. Numbers represent percent cell marker normalized to DAPI, mean ± SEM. (**B**) Immunostaining of iPSN cultures for key neuronal proteins. Genotype as indicated on left, time point and antibodies as indicated on top. (**C**) qRT-PCR for G_4_C_2_ and C_2_G_4_ repeat RNA in *C9orf72* iPSNs. (**D**) MSD Elisa for Poly(GP) DPR levels. Shapes indicate individual C9orf72 iPSC lines and corresponding black colored shape indicates paired control iPSC line. (**E**) qRT-PCR for total *C9orf72* RNA. (**F-G**) Western blot (**F**) and quantification (**G**) for C9ORF72 protein. (**H**) Confocal imaging of iPSNs immunostained for TDP-43. Genotype as indicated on left, antibodies as indicated on top. (**I**) Quantification of nuclear to cytoplasmic ratio of TDP-43. N = 4 control and 4 *C9orf72* iPSC lines, 50 cells per line. Student’s T-test was used to calculate statistical significance. (**J**) Percent of cell death following exposure to glutamate. N = 3 control and 3 C9orf72 iPSC lines, 10 frames per well. Two-way ANOVA with Tukey’s multiple comparison test was used to calculate statistical significance. *** p < 0.001, **** p < 0.0001. Scale Bar = 500 μm (**B**) 10 μm (**H**).

**Figure S2.**
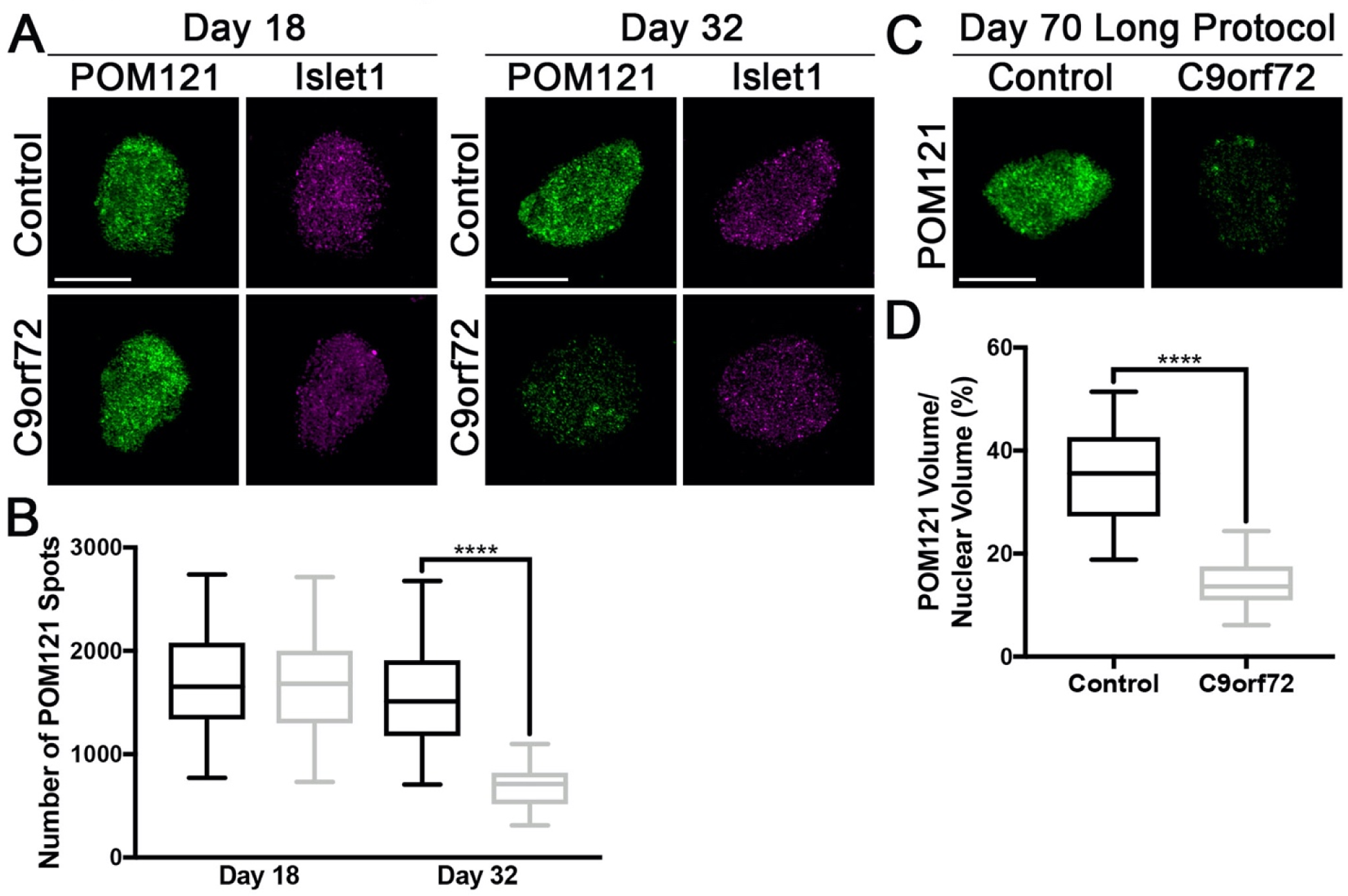
Related to Figure 1: POM121 is reduced in Islet1 positive nuclei and nuclei from multiple spinal neuron differentiation protocols. (**A-B**) Maximum intensity projections from SIM imaging (**A**) and quantification (**B**) of POM121 spots in Islet1+ nuclei isolated from control and *C9orf72* iPSNs. Genotype as indicated on left, time point and antibody as indicated on top. N = 4 control and 4 *C9orf72* iPSC lines (including 1 isogenic pair), 50 Islet1+ nuclei per line/time point. Two-way ANOVA with Tukey’s multiple comparison test was used to calculate statistical significance. **** p < 0.0001. (**C-D**) SIM imaging (**C**) and quantification (**D**) of POM121 in day 70 nuclei isolated from control and *C9orf72* iPSNs differentiated using a longer spinal neuron differentiation protocol. Antibody as indicated on left, genotype as indicated on top. N = 3 control and 3 *C9orf72* iPSC lines, 50 NeuN+ nuclei per line. Student’s T-test was used to calculate statistical significance. **** p < 0.0001. Scale bar = 10 μm.

**Figure S3.**
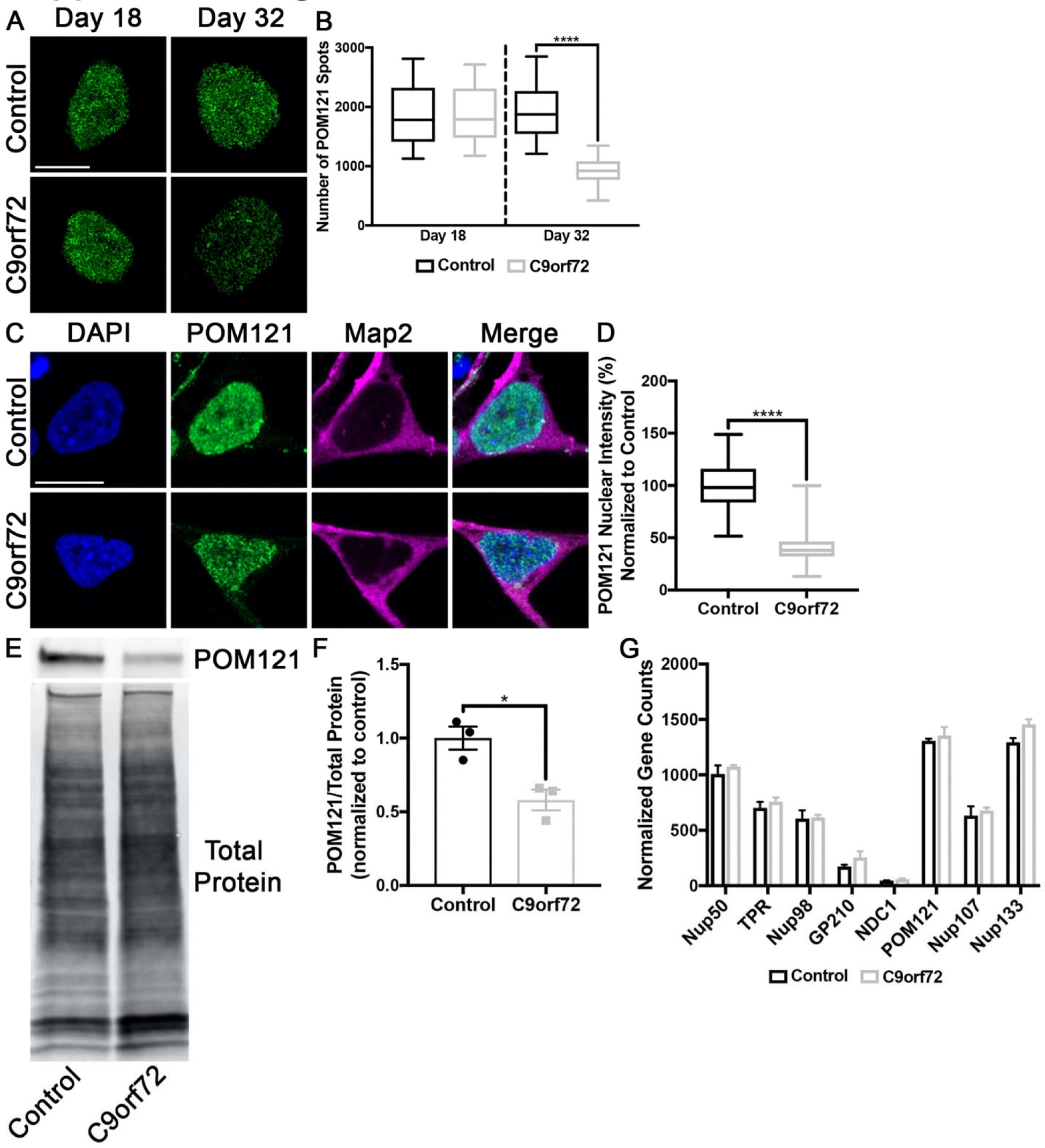
Related to Figure 1: POM121 protein is reduced in *C9orf72* iPSNs. (**A**) Maximum intensity projections from SIM imaging of POM121 in nuclei isolated from control and *C9orf72* iPSNs. Genotype as indicated on left, time point as indicated on top. (**B**) Quantification of POM121 spots. N = 8 control and 8 *C9orf72* iPSC lines (including 1 isogenic pair), 50 NeuN+ nuclei per line/time point. Two-way ANOVA with Tukey’s multiple comparison test was used to calculate statistical significance. **** p < 0.0001. (**C**) Confocal imaging of control and *C9orf72* iPSNs immunostained for POM121. Genotype as indicated on left, antibodies as indicated on top. (**D**) Quantification of POM121 nuclear intensity. N = 3 control and 3 *C9orf72* iPSC lines, at least 50 cells per line. Student’s T-test was used to calculate statistical significance. **** p < 0.0001. (**E-F**) Western blot (**E**) and quantification (**F**) for POM121 from nuclei isolated from control and *C9orf72* iPSNs. Antibodies as indicated on right, genotype as indicated on bottom. N = 3 control and 3 *C9orf72* iPSC lines. Student’s T-test was used to calculate statistical significance. * p < 0.05. (**G**) Nanostring gene expression analysis for Nups whose NPC localization and expression is altered in *C9orf72* iPSNs. Scale bar = 5 μm (**A**), 10 μm (**C**).

**Figure S4.**
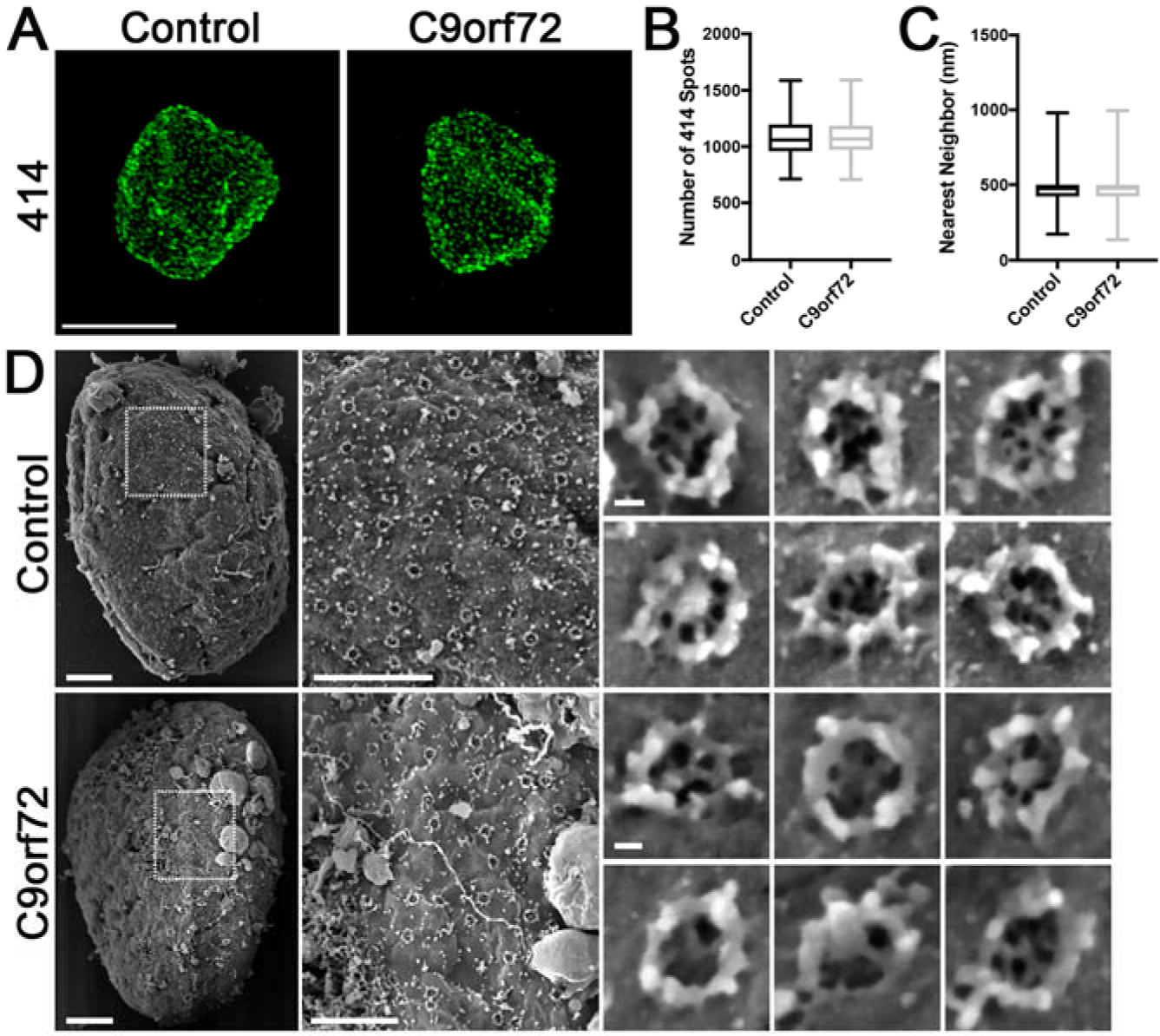
Related to Figure 1: Overall NPC number, distribution, and structure is unchanged in *C9orf72* iPSNs. (**A**) Maximum intensity projections from SIM imaging of 414 in nuclei isolated from control and *C9orf72* iPSNs. Genotype as indicated on top. (**B-C**) Quantification of NPC spots (**C**) and distribution (**D**) as determined by 414 staining. N = 8 control and 8 *C9orf72* iPSC lines (including 1 isogenic pair), 50 NeuN+ nuclei per line. Student’s T-test was used to calculate statistical significance. (**D**) Direct surface imaging of NPCs by scanning electron microscopy on nuclei from control and *C9orf72* iPSNs. Genotype as indicated on left. Scale bar = 5 μm (**A**), from left to right 1 μm, 500 nm, 20 nm (**D**).

**Figure S5.**
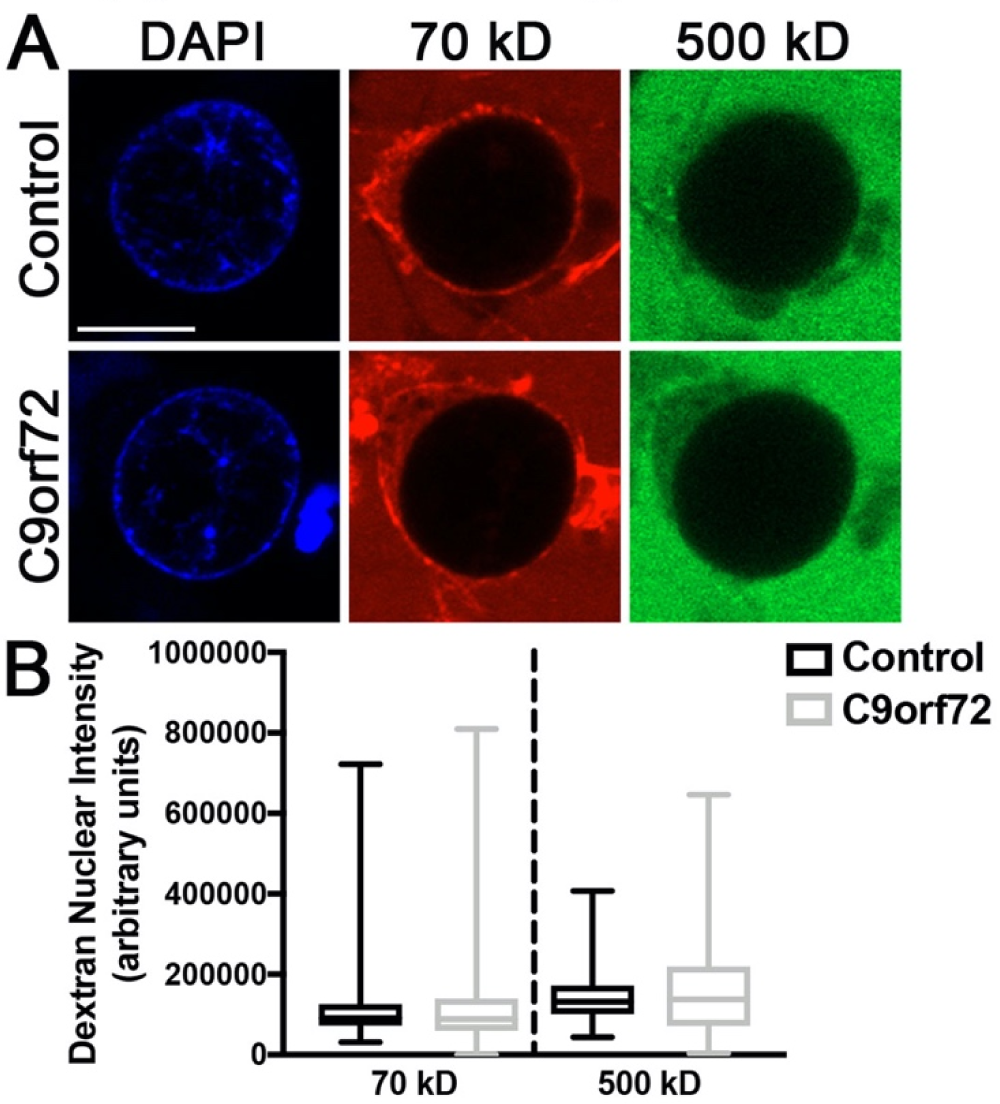
Related to Figure 1: *C9orf72* iPSN nuclei are not leaky. (**A**) Confocal imaging of fluorescently tagged dextrans in digitonin permeabilized control and *C9orf72* iPSNs. Genotype as indicated on left, dextran as indicated on top. (**B**) Quantification of nuclear intensity of dextrans. N = 6 control and 6 *C9orf72* iPSC lines, at least 50 nuclei per line. Two-way ANOVA with Tukey’s multiple comparison test was used to calculate statistical significance. Scale bar = 10 μm.

**Figure S6.**
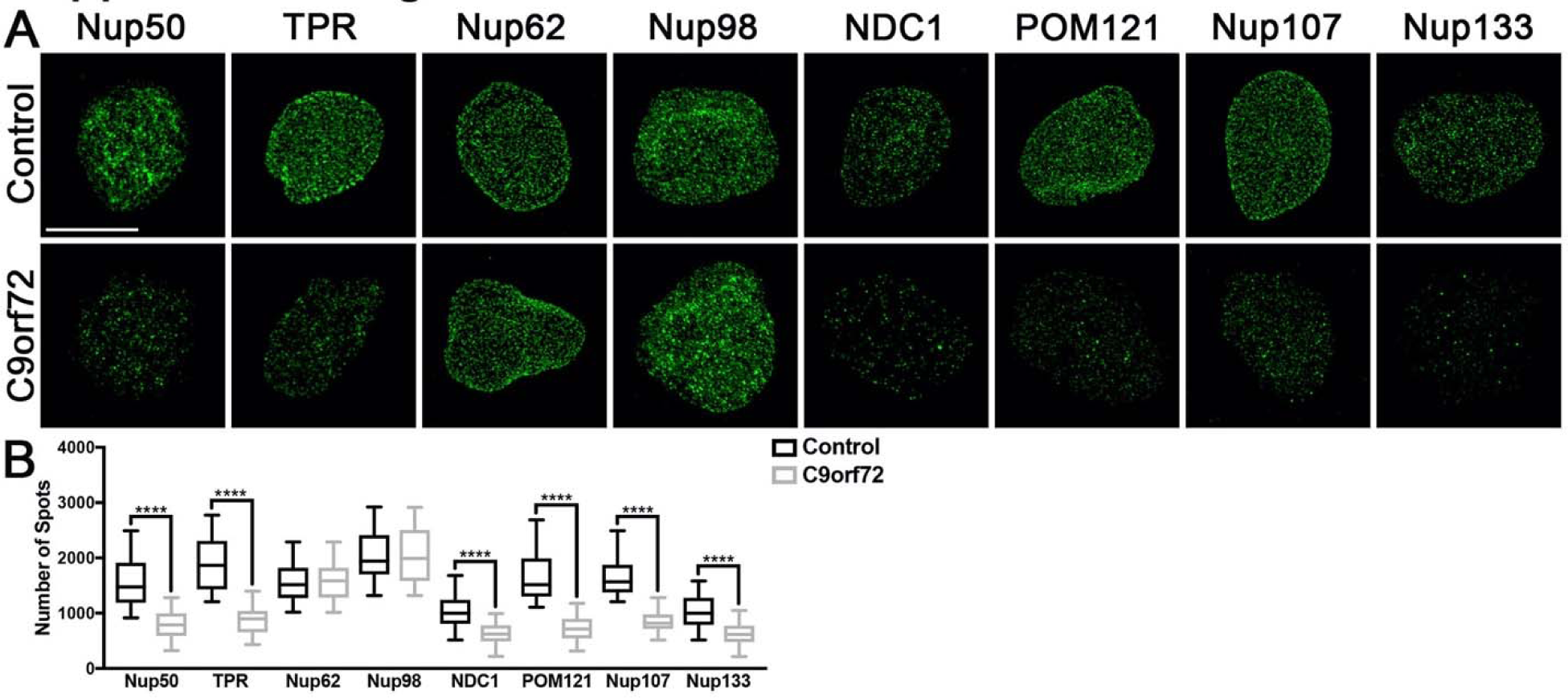
Related to Figure 2: Specific nucleoporin alterations are recapitulated in nuclei from postmortem C9orf72 thoracic spinal cord. (**A**) Maximum intensity projections from SIM imaging of Nups in nuclei isolated from postmortem human thoracic spinal cord. Genotype as indicated on left, antibodies as indicated on top. (**B**) Quantification of Nup spots and volume. N = 3 control and 3 *C9orf72* cases, 50 NeuN+ nuclei per case. Two-way ANOVA with Tukey’s multiple comparison test was used to calculate statistical significance. **** p < 0.0001. Scale bar = 5 μm.

**Figure S7.**
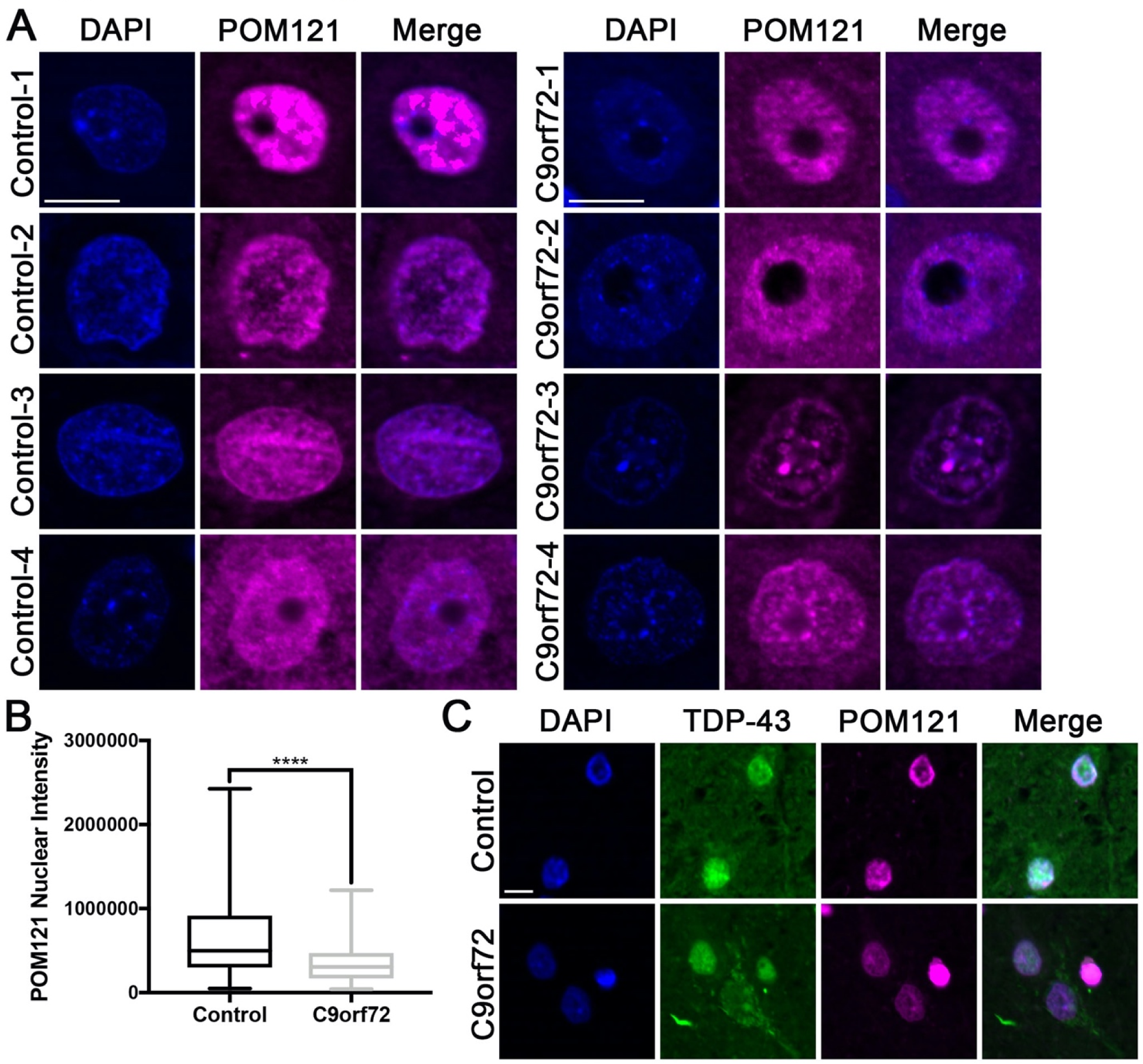
Related to Figure 2: POM121 is reduced but not mislocalized in postmortem *C9orf72* patient motor cortex. (**A**) Immunostaining for POM121 in paraffin embedded postmortem motor cortex. (**B**) Quantification of POM121 nuclear intensity. N = 4 control and 4 *C9orf72* cases, at least 100 nuclei from Map2+ cells per case. Student’s T-test was used to calculate statistical significance. **** p < 0.0001. (**C**) Immunostaining for TDP-43 and POM121 in paraffin embedded postmortem motor cortex. Genotype as indicated on left, antibodies as indicated on top. Scale bar = 10 μm.

**Figure S8.**
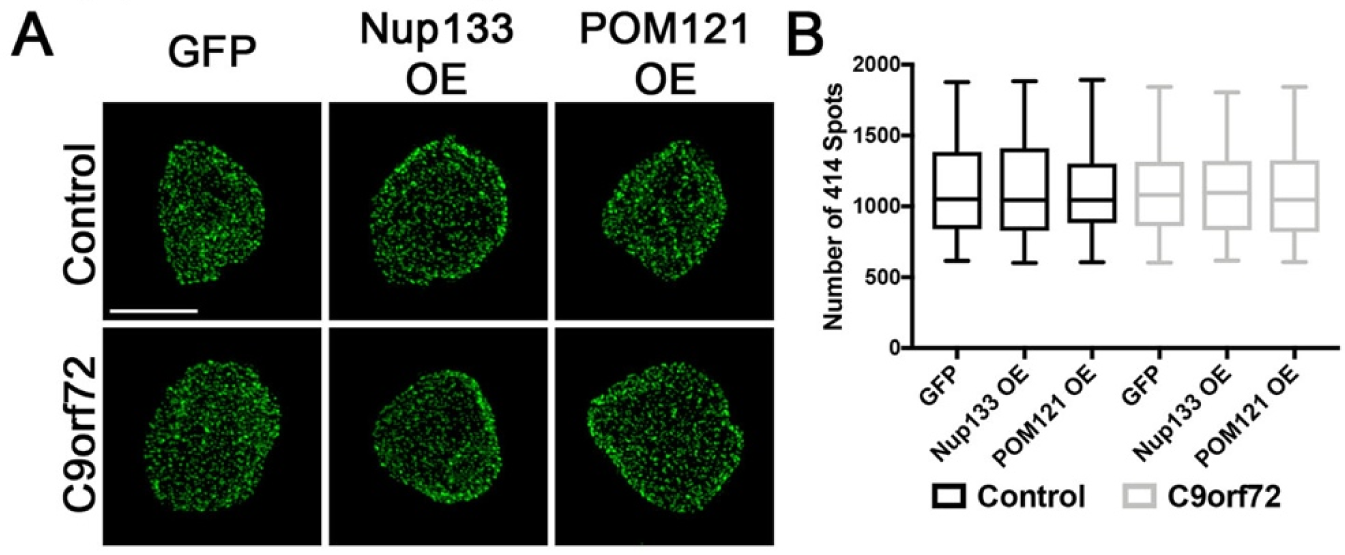
Related to Figure 3: Overexpression of POM121 or Nup133 does not alter the total number of nuclear pores. (**A**) Maximum intensity projections from SIM imaging of 414 spots in nuclei isolated from control and *C9orf72* iPSNs overexpressing Nup133 or POM121. Genotype as indicated on left, overexpression as indicated on top. (**B**) Quantification of 414 spots. N = 4 control and 4 *C9orf72* iPSC lines, 50 GFP+ nuclei per line/overexpression. Two-way ANOVA with Tukey’s multiple comparison test was used to calculate statistical significance.

**Figure S9.**
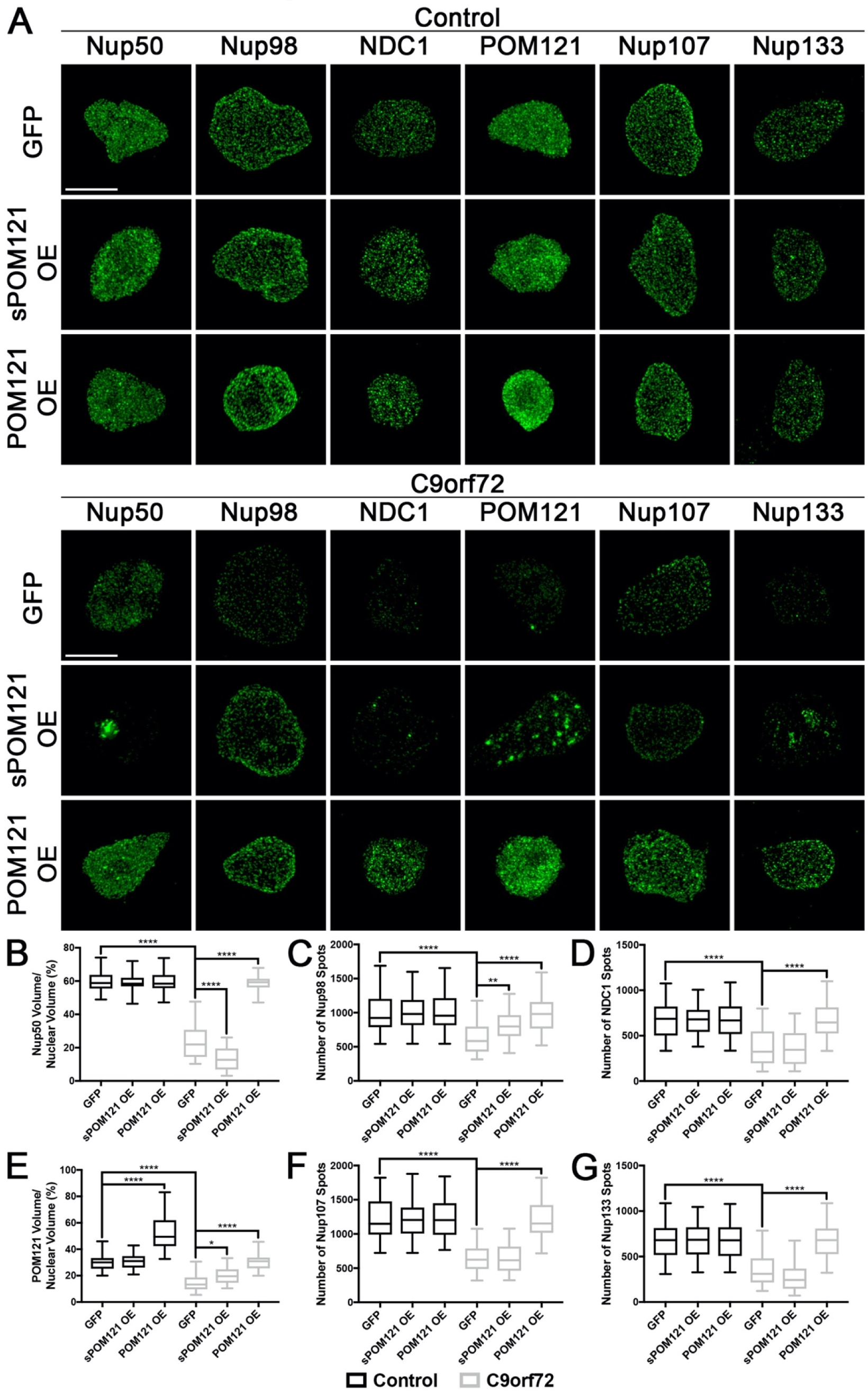
Related to Figure 3: sPOM121 does not contribute to restored nuclear expression and localization of Nups in *C9orf72* iPSNs. (**A**) Maximum intensity projections from SIM imaging of Nups in nuclei isolated from control and *C9orf72* iPSNs overexpressing sPOM121 or POM121. Overexpression as indicated on left, genotype and antibodies as indicated on top. (**B-G**) Quantification of Nup spots and volume. N = 3 control and 3 *C9orf72* iPSC lines, 50 GFP+ nuclei per line/overexpression. Two-way ANOVA with Tukey’s multiple comparison test was used to calculate statistical significance. * p < 0.05, ** p < 0.01, **** p < 0.0001.

**Figure S10.**
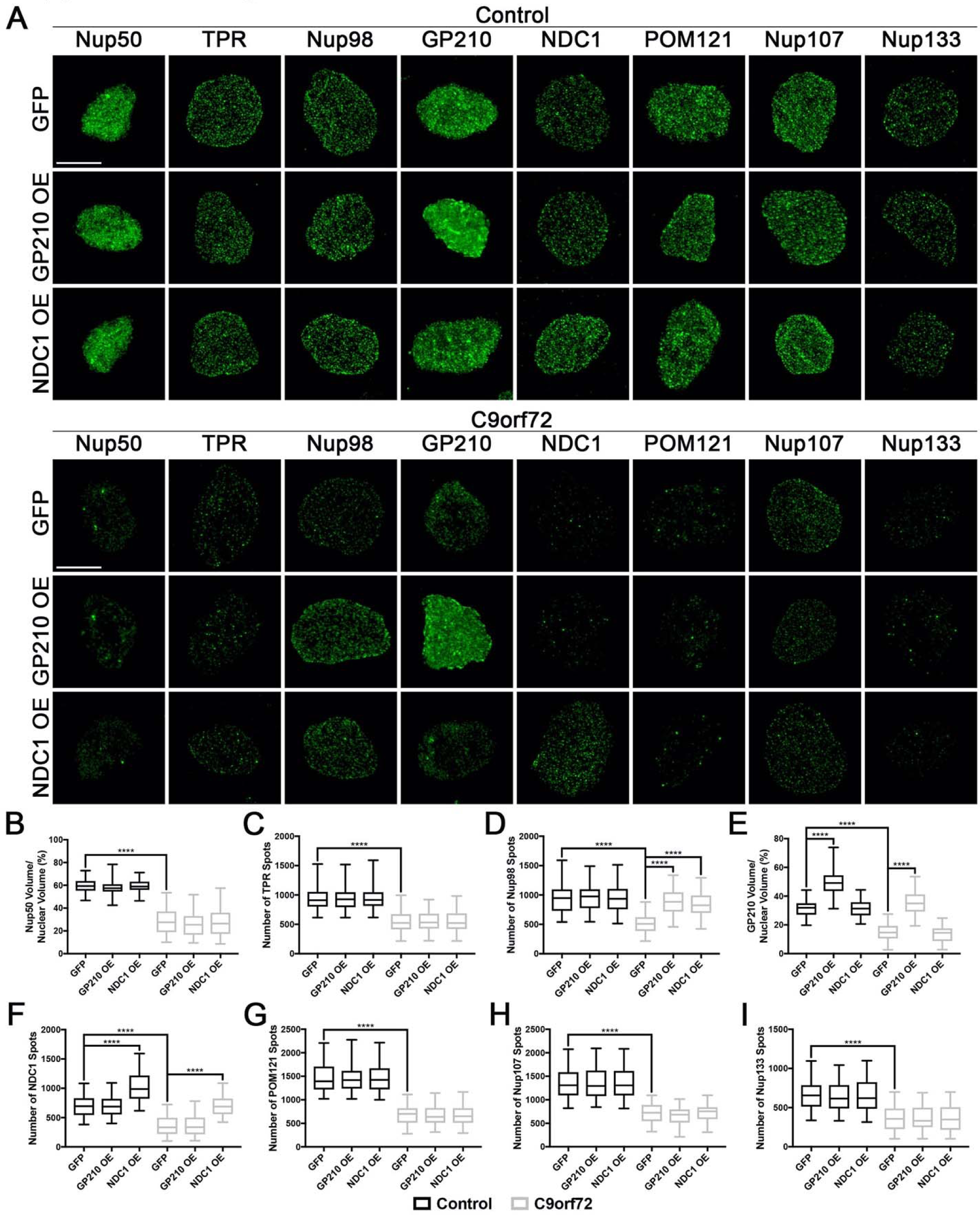
Related to Figure 3: Overexpression of the transmembrane Nups GP210 or NDC1 does not contribute to restored nuclear expression and localization of affected Nups in *C9orf72* iPSNs. (**A**) Maximum intensity projections from SIM imaging of Nups in nuclei isolated from control and *C9orf72* iPSNs overexpressing GP210 or NDC1. Overexpression as indicated on left, genotype and antibodies as indicated on top. (**B-I**) Quantification of Nup spots and volume. N = 3 control and 3 *C9orf72* iPSC lines, 50 GFP+ nuclei per line/overexpression. Two-way ANOVA with Tukey’s multiple comparison test was used to calculate statistical significance. **** p < 0.0001. Scale bar = 5 μm.

**Figure S11.**
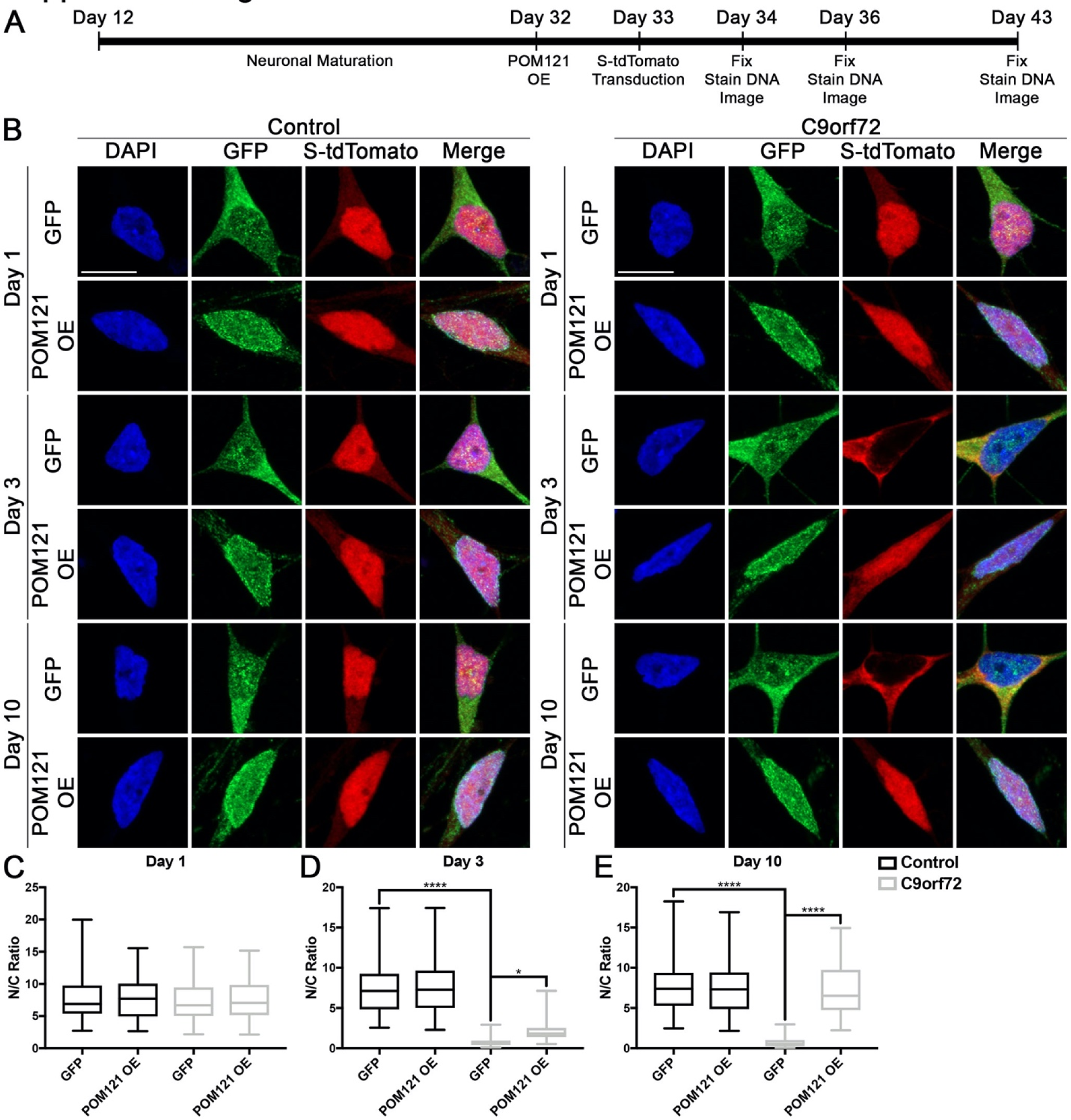
Related to Figure 3: POM121 overexpression restores the localization of the S-tdTomato reporter in *C9orf72* iPSNs. (**A**) Schematic representation of experimental time line. (**B**) Confocal imaging of control and *C9orf72* iPSNs overexpressing POM121 GFP and the NLS-tdTomato-NES (S-tdTomato) NCT reporter construct. Time point and overexpression as indicated on left, genotype and reporter as indicated on top. (**C-E**) Quantification of nuclear to cytoplasmic ratio of the S-tdTomato reporter at day 1 (**C**), day 3 (**D**), and day 10 (**E**) post transduction. N = 3 control and 3 *C9orf72* iPSC lines, at least 25 cells per line/overexpression/time point. Two-way ANOVA with Tukey’s multiple comparison test was used to calculate statistical significance. * p < 0.05, **** p < 0.0001. Scale bar = 10 μm.

**Figure S12.**
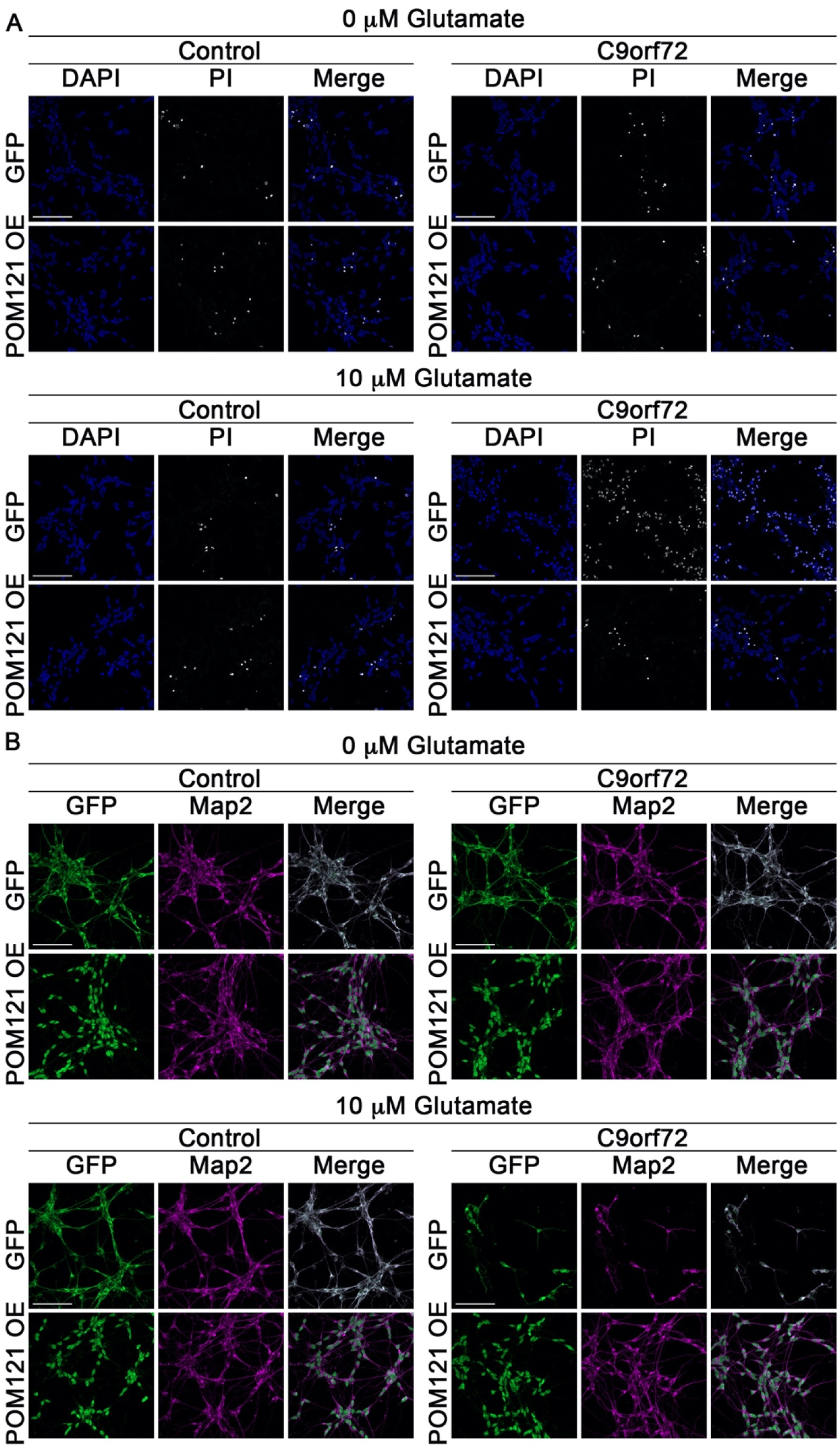
Related to Figure 3: POM121 overexpression mitigates glutamate induced excitotoxicity in *C9orf72* iPSNs. (**A**) Confocal imaging of cell death in control and C9orf72 iPSNs as measured by propidium iodide (PI) incorporation. Overexpression as indicated on left, genotype and stain as indicated on top. (**B**) Confocal imaging of surviving neurons immunostained for POM121 GFP and Map2 following exposure to glutamate. Overexpression as indicated on left, genotype and antibodies as indicated on top. N = 4 control and 4 *C9orf72* iPSC lines, 10 frames per well. Scale bar = 100 μm.

**Figure S13.**
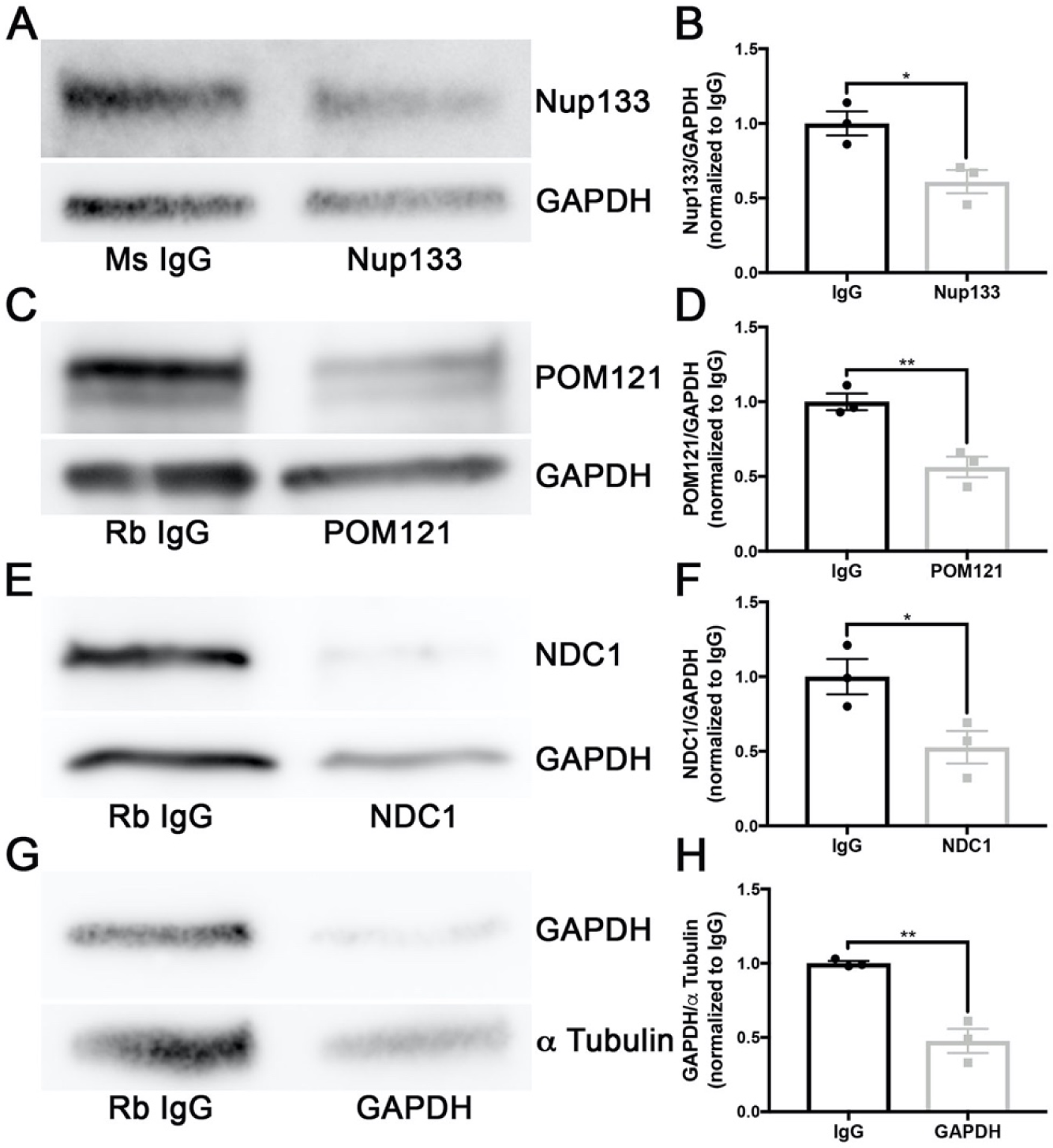
Related to Figure 4: Trim21 mediated knock down of nucleoporins and GAPDH in wildtype iPSNs. Western blots and quantification of Trim21 GFP mediated reduction in POM121 (**A-B**), Nup133 (**C-D**), NDC1 (**E-F**), and GAPDH (**G-H**) in wildtype iPSNs. Student’s T-test was used to calculate statistical significance. * p < 0.05, ** p < 0.01.

**Figure S14.**
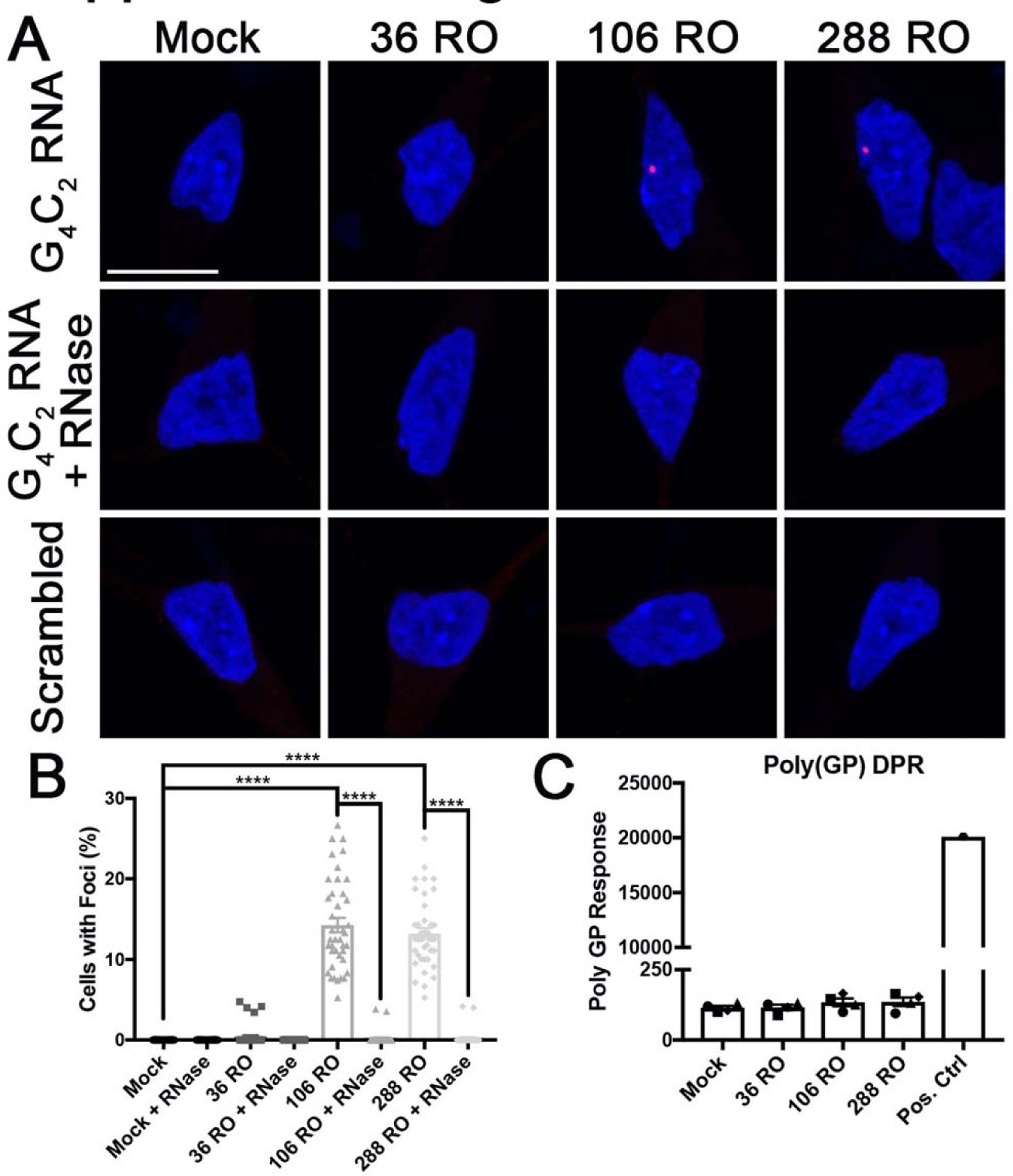
Related to Figure 6: G_4_C_2_ repeat RNA only constructs form RNA foci but do not produce DPRs in wildtype iPSNs. (**A**) Confocal imaging of RNA FISH for G_4_C_2_ repeat RNA foci in wildtype iPSNs overexpressing G_4_C_2_ repeat RNA only constructs. Probe and treatment as indicated on left, overexpression as indicated on top. (**B**) Quantification of the percent of cells containing G_4_C_2_ RNA foci. N = 4 control iPSC lines, at least 150 cells across 10 frames per line/overexpression. Two-way ANOVA with Tukey’s multiple comparison test was used to calculate statistical significance. **** p < 0.0001. Individual data points represent each image frame. (**C**) MSD Elisa for Poly(GP) DPR expression in wildtype iPSNs overexpressing G_4_C_2_ RNA only constructs. Shapes indicate individual C9orf72 iPSC lines. Scale bar = 10 μm.

**Figure S15.**
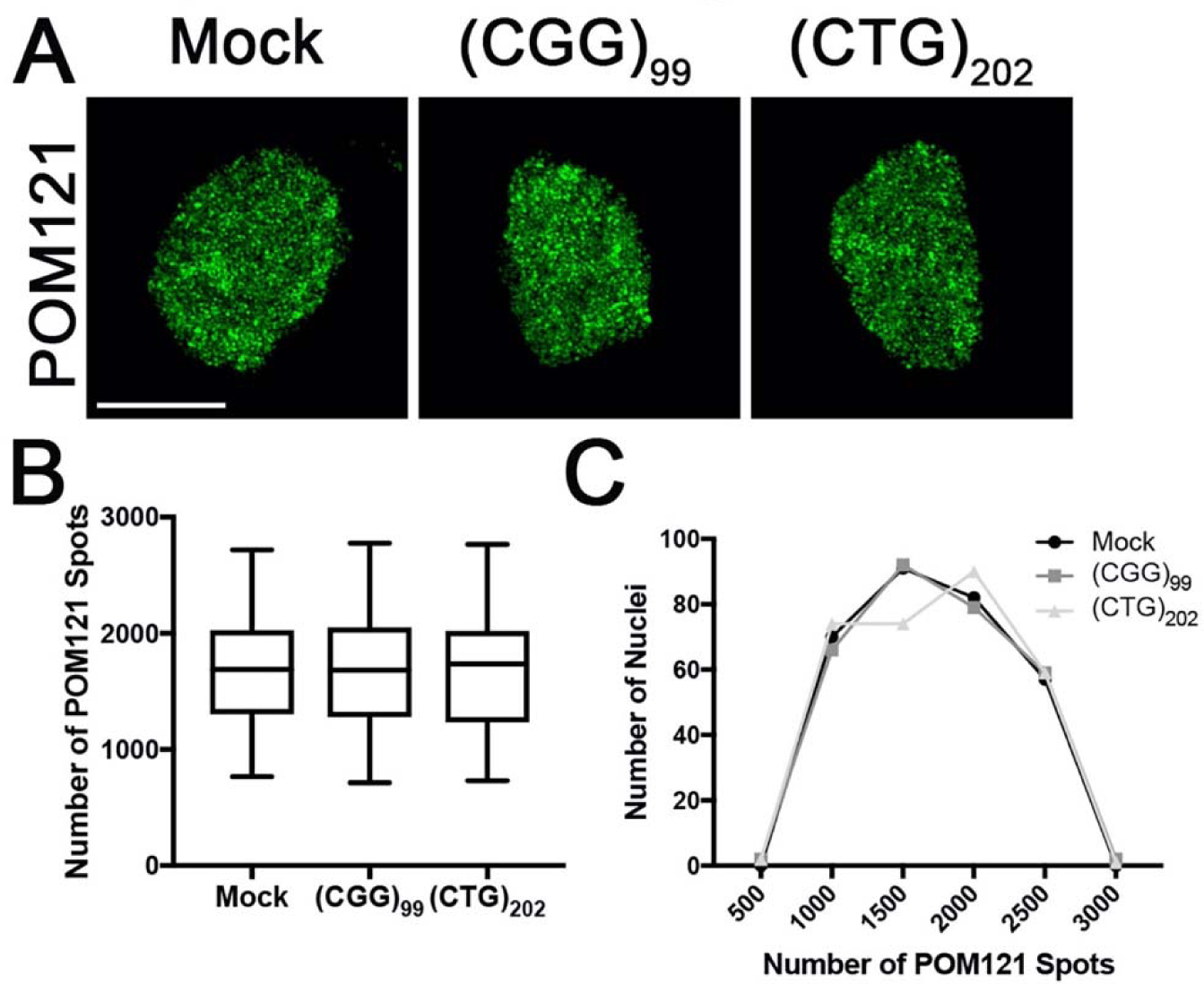
Related to Figure 6: Overexpression of CGG and CTG repeat RNAs does not affect the nuclear localization and expression of POM121 in iPSNs. (**A**) Maximum intensity projections from SIM imaging of POM121 in nuclei isolated from wildtype iPSNs overexpressing (CGG)_99_ or (CTG)_202_ repeat RNAs for 7 days. Antibody as indicated on left, overexpression as indicated on top. (**B-C**) Quantification and histogram distributions of POM121 spots 7 days after overexpression of repeat RNAs. N = 3 wildtype iPSC lines, 100 NeuN+ nuclei per line/overexpression. One-way ANOVA with Tukey’s multiple comparison test was used to calculate statistical significance. **** p < 0.0001. Scale bar = 10 μm.

**Figure S16.**
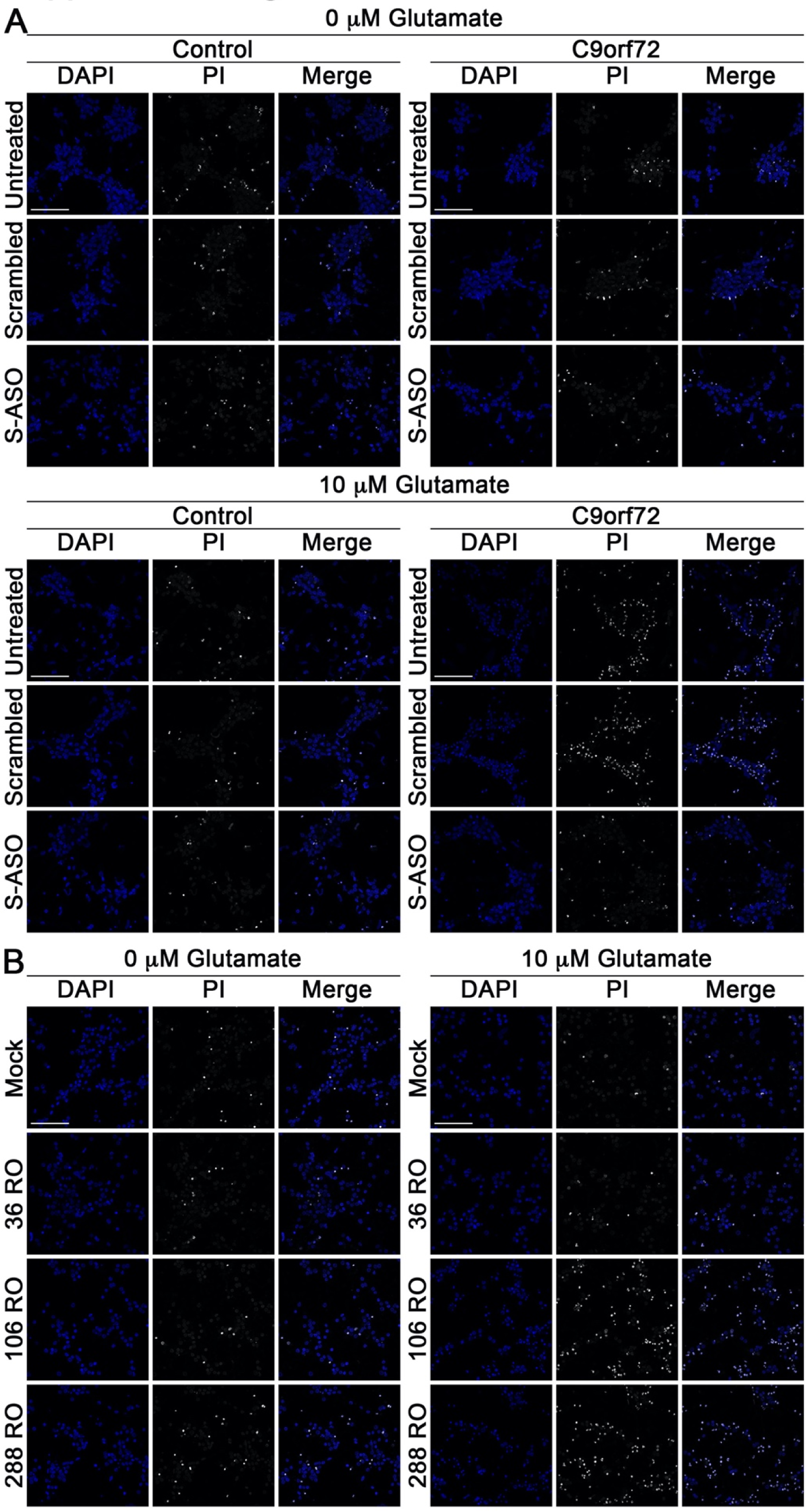
Related to Figure 6: Expression of pathological G_4_C_2_ repeat RNA renders iPSNs susceptible to glutamate induced excitotoxicity. (**A**) Confocal imaging of cell death in control and C9orf72 iPSNs following 5 day exposure to G_4_C_2_ targeting ASO as measured by propidium iodide (PI) incorporation. Treatment as indicated on left, genotype and stain as indicated on top. N = 4 control and 4 *C9orf72* iPSC lines, 10 frames per well. (**B**) Confocal imaging of cell death in wildtype iPSNs overexpressing G_4_C_2_ repeat RNA as measured by propidium iodide (PI) incorporation. Overexpression as indicated on left, genotype and stain as indicated on top. N = 4 control iPSC lines, 10 frames per well. Scale bar = 100 μm.

**Figure S17:**
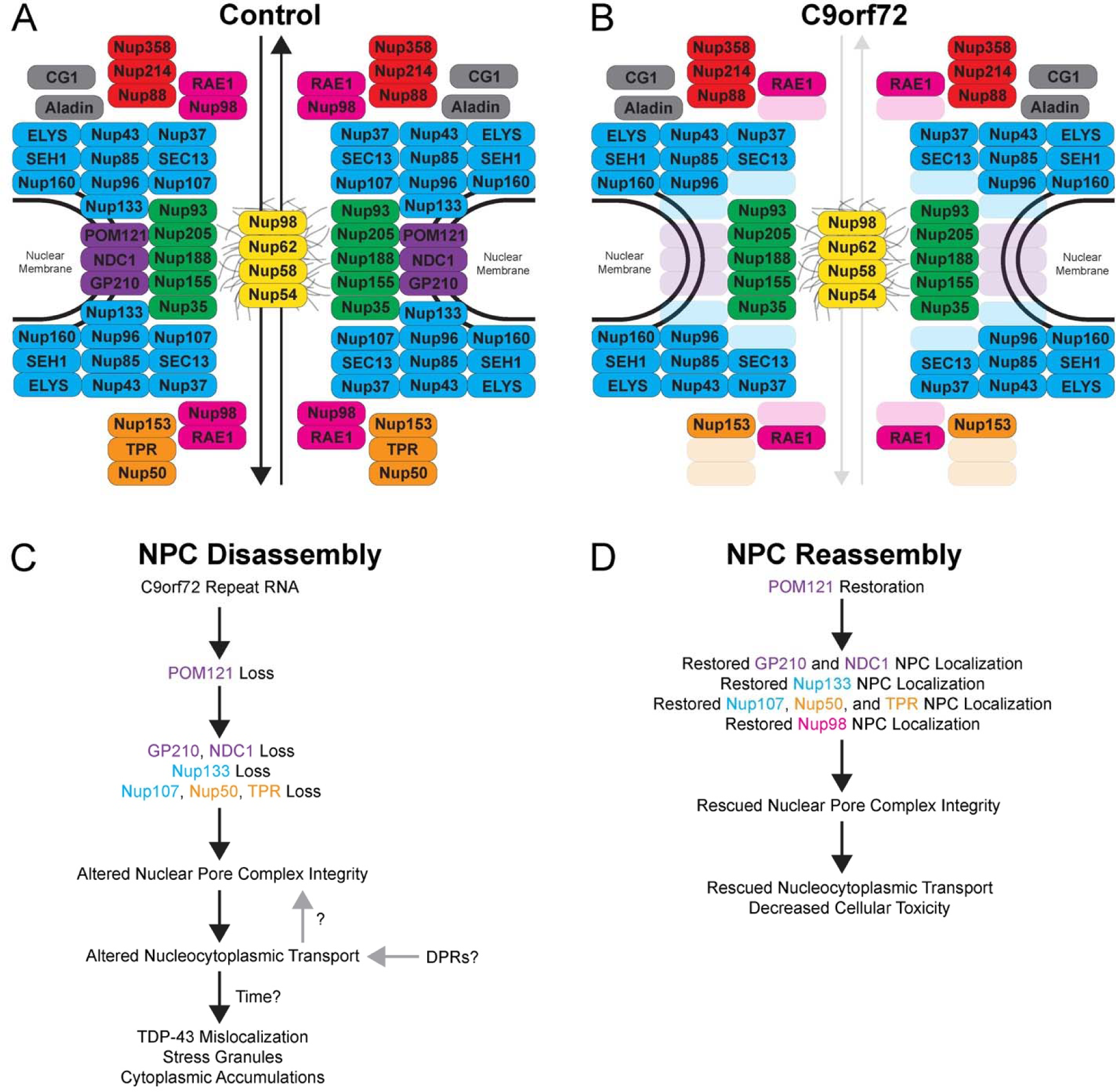
A model for NPC assembly and disassembly in *C9orf72* ALS. (**A-B**) The structure of the NPC in normal (**A**) and disease (**B**) neurons. (**C**) NPC disassembly is initiated by direct or indirect *C9orf72* HRE mediated loss of POM121. Loss of POM121 results in the disassembly of the NPC beginning with the remainder of the transmembrane Nups (GP210, NDC1), followed by the Y complex outer ring Nups (Nup133, Nup107) and the nuclear basket Nups (Nup50, TPR) ultimately leading to disrupted NCT. (**D**) Neuronal NPC reassembly is initiated by integration of POM121 into the NPC. Restored NPC integrity rescues NCT and decreases susceptibility to glutamate induced excitotoxicity.

